# Wilson disease: intersecting DNA methylation and histone acetylation regulation of gene expression in a mouse model of hepatic copper accumulation

**DOI:** 10.1101/2021.04.15.439900

**Authors:** Gaurav V. Sarode, Kari Neier, Noreene M. Shibata, Yuanjun Shen, Dmitry A Goncharov, Elena A. Goncharova, Tagreed A. Mazi, Nikhil Joshi, Matthew L. Settles, Janine M. LaSalle, Valentina Medici

## Abstract

The pathogenesis of Wilson disease (WD) is multi-factorial, involving hepatic and brain copper accumulation due to pathogenic variants affecting the *ATP7B* gene and downstream epigenetic and metabolic mechanisms. Prior DNA methylation investigations in human WD liver and blood and in a WD mouse model revealed an epigenetic signature of WD, including alterations in the histone deacetylase HDAC5. To test the hypothesis that histone acetylation is altered with respect to copper overload and aberrant DNA methylation in WD, we investigated class IIa histone deacetylases (HDAC4 and HDAC5) and H3K9/H3K27 histone acetylation in the Jackson Laboratory toxic milk (tx-j) mouse model of WD compared to C3HeB/FeJ (C3H) control in response to 3 treatments: 60% kcal fat diet (HFD), D-penicillamine (PCA, copper chelator), and choline (methyl group donor). HDAC5 levels significantly increased in 9-week tx-j livers after 8 days of HFD compared to chow. In 24-week tx-j livers, HDAC4/5 levels were reduced 5- to 10-fold compared to C3H likely through mechanisms involving HDAC phosphorylation. HDAC4/5 levels were also affected by disease progression and accompanied by increased acetylation. PCA and choline partially restored HDAC4, HDAC5, H3K9ac, and H3K27ac levels to that of CH3 liver. Integrated RNA and chromatin immunoprecipitation sequencing analyses revealed genes regulating energy metabolism and cellular stress/development were, in turn, regulated by histone acetylation in tx-j mice compared to C3H, with *Pparα* and *Pparγ* among the most relevant targets. These results suggest dietary modulation of class IIa HDAC4/5, and subsequent H3K9/H3K27 acetylation/deacetylation, can regulate gene expression in key metabolic pathways in the pathogenesis of WD.

**Significance Statement:** Wilson disease is considered a monogenic disease caused by pathogenic variants in the ATP7B copper transporter, resulting in hepatic and brain copper accumulation. Given the lack of genotype-phenotype correlation, evidence of epigenetic and metabolic mechanisms regulating phenotype in patients and in animal models could explain the high phenotype variability observed in WD. In this study, we identify class IIa histone deacetylases as players involved in the epigenetic regulation of key metabolic pathways that can affect WD severity as well as targets sensitive to dietary modulations, which is an important characteristic for designing effective and feasible therapies. Understanding the epigenetic mechanisms in WD pathogenesis contributes to a better understanding of the phenotypic variability in WD and other common liver conditions.

## Introduction

Wilson disease (WD) is a systemic disease caused by the inheritance of recessive mutations affecting the *ATP7B* copper transporter gene. As a result of transporter loss of function, copper accumulates predominantly in the liver and the brain. Although hepatic and neurologic signs and symptoms are the most common manifestations, any organ can be affected with various degrees of involvement. The growing knowledge about WD pathogenesis has also contributed to the understanding of more common conditions, including alcohol-associated liver disease, nonalcoholic fatty liver, and autoimmune hepatitis (1-3), as the involved pathogenic pathways partially overlap and the clinical and laboratory presentations have common features (4-6). Although the genetic deficiency in WD causes copper accumulation, gene variants themselves do not explain the whole array of clinical manifestations as there is no evidence of genotype-phenotype correlation (7). Given the similarity of systemic involvement and hepatic phenotypes with other metabolic liver diseases, WD is ideal for understanding the mechanisms leading to epigenetic and metabolic dysregulation during disease onset and progression as well as the extent of organ involvement. Moreover, genetic mouse models of hepatic copper accumulation replicate many of the metabolic derangements observed in humans (8-11).

One-carbon metabolism and DNA methylation are among the processes identified as being aberrant in WD, with specific patterns distinguishing WD from other liver diseases (12, 13). The mechanism underlying the connection between copper accumulation and methionine metabolism involves an inhibitory effect of copper on the enzyme S-adenosylhomocysteine (SAH) hydrolase with consequent accumulation of SAH; SAH can interfere with DNA methylation mechanisms by inhibiting DNA methyltransferases. DNA methylation alterations are observed in patients with WD and in the Jackson Laboratory toxic milk mouse C3He-Atp7b^tx-j^/J (tx-j), a model of spontaneous hepatic copper accumulation (9). Hypermethylated differentially methylated regions were enriched in enhancer sites, suggesting functional relevance of methylation changes affecting gene expression regulation. DNA methylation changes partially overlapped between patients and mice; changes were evident in fetal and adult mouse livers (10) as well as in human blood and liver, indicating a universal presence of DNA methylation alterations characterizing WD. Furthermore, mechanistically meaningful pathways were identified from DNA methylation analyses that could lead to a deeper understanding of WD pathogenesis. Interestingly, among the identified regulatory genes, histone deacetylase 5 (*HDAC5*) was hypermethylated in the gene body both in blood and liver tissue from patients with WD compared to healthy subjects. HDACs, which are categorized into classes I to IV, catalyze histone deacetylations, resulting in epigenetic chromatin modifications that inhibit gene expression. HDAC5 is a class IIa enzyme shuttled between the nucleus and cytosol when phosphorylated in a rapid response to various environmental stimuli, including nutrients, inflammation, and oxidative stress (14). Therefore, in addition to DNA methylation, HDACs can be considered an interface between nutrition and gene expression regulation. Increasing evidence from cancer literature indicates a multi-layered interplay between chromatin structure, histone acetylation, and DNA methylation. Inhibition of DNA methyltransferases or HDACs is known to have synergistic effects on cellular replication and differentiation, as observed in *in vitro* cancer models (15, 16). Histone post-translational modifications, including trimethylation of histone-3-lysine sites (H3K9me3 and H3K27me3), are commonly known to be associated with regulatory loci.

There is growing interest regarding the role of epigenetic mechanisms in metabolic conditions whose presentation and liver involvement can mimic WD. Hepatic class I and II HDAC transcript levels and activities are affected by ethanol consumption in a mouse model of alcoholic liver disease and HDAC levels are dysregulated at the promoter of genes critical for liver damage progression (17). Alcohol-induced histone 3 acetylation has been extensively studied in alcoholic liver disease and it appears to play a role in the progression of fibrosis (18, 19). HDAC5 changes have also been implicated in liver damage when C57Bl/6 mice were fed a high fat diet (20). Acetylated H3K9 (H3K9ac) and 5-methylcytosine levels were altered at regulatory loci of aging-associated genes in mouse livers (21). There is, however, lack of knowledge in WD about the network involving methyl group availability and its effects on histone acetylation. This is crucial information as in-depth understanding of epigenetic mechanisms and their cross-talk with metabolic factors and gene expression regulation will contribute to a better understanding of WD phenotypic variability as well as the metabolic underpinnings of more common liver conditions with similar hepatic presentations. In this study, we tested the hypothesis that hepatic copper levels and methyl group availability interact with HDAC5 and histone acetylation in regulating gene transcript levels in animal models of WD.

## Results

### Histone deacetylase 5 is downregulated in multiple models of Wilson disease

To test the hypothesis that epigenetic changes to *HDAC5* resulted in changes to protein levels, Western blot analyses were performed in multiple *in vivo* and *in vitro* models of WD (see animal study design Figure S1). First, HDAC5 protein levels were reduced in tx-j livers compared to C3HeB/FeJ (C3H) control mice (Figure 1A, S2A). At 6 days post-partum, when hepatic copper levels are still normal, hepatic HDAC5 levels were about 1.3-fold lower in tx-j mice. The difference became progressively more significant, concomitantly with copper accumulation. Hepatic HDAC5 levels were about 5-fold lower in 12-week old tx-j mice to 10-fold lower in 24-week old tx-j mice compared to C3H. Similarly, class IIa histone deacetylase HDAC4 was also reduced in tx-j mouse livers, indicating a widespread effect on class II HDACs. HDAC4 showed reduced expression at the latest stages of disease progression from 12-to 24-weeks. In the *Atp7b*^-/-^mouse model of WD, HDAC4 and HDAC5 levels were also significantly reduced in 16-week old *Atp7b*^-/-^compared to wild-type mice (Figure 1B, S2B).

**Figure 1.**
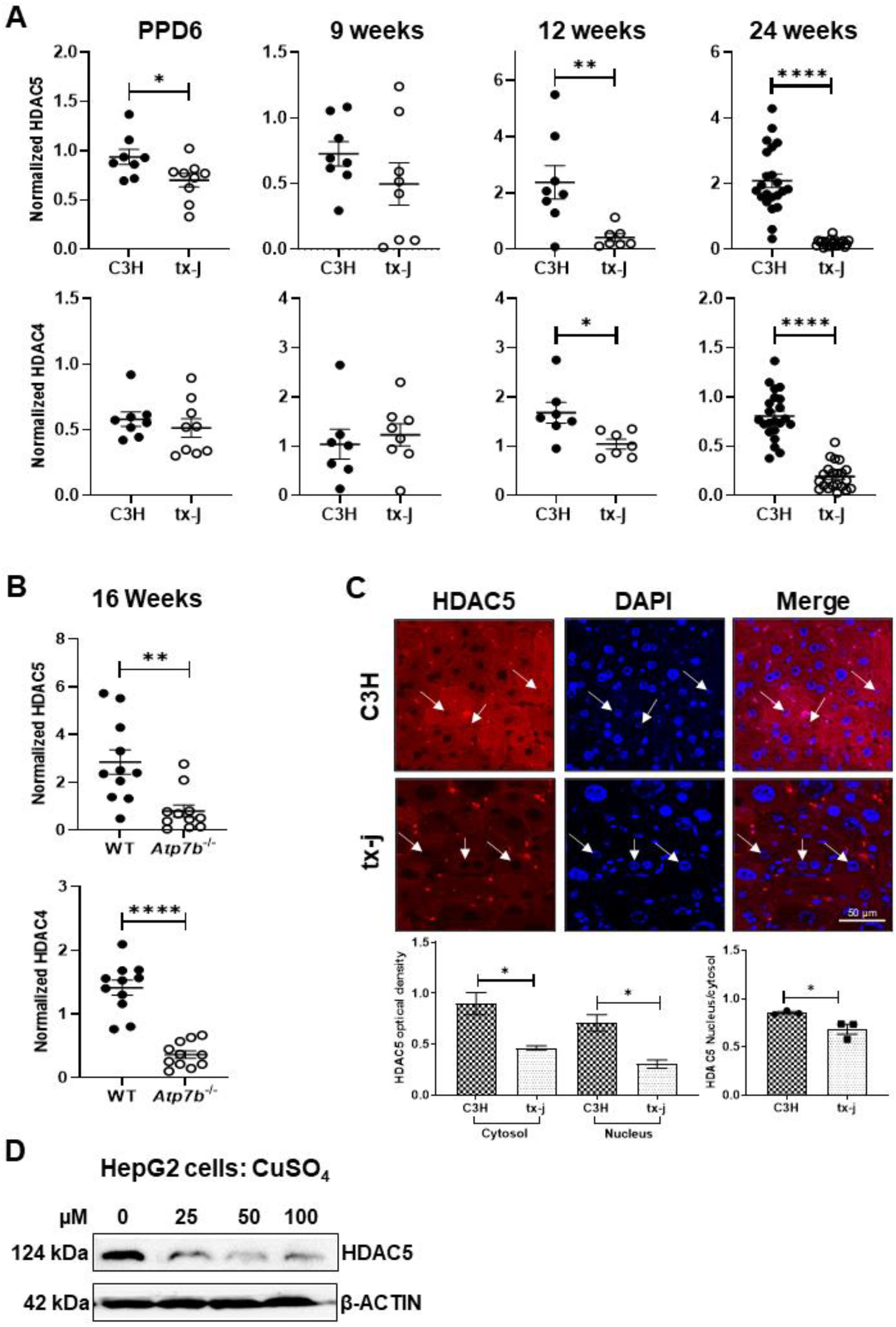
Expression of HDAC4 and HDAC5 in mouse and HepG2 models of WD. Immunoblot densitometry analyses are normalized to β-actin; data are represented as means ± SEM and statistical significance was determined by Student’s t-test (*p< 0.05, **p< 0.01, and ****p< 0.0001). A: Total protein liver lysate densitometries for HDAC4 and HDAC5 protein expression in tx-j mice compared to C3H control at post-partum day 6 (PPD6; C3H n=4M/4F, tx-j n=5M/4F), 9 weeks (C3H n=4M/3F, tx-j n=4M/4F), 12 weeks (C3H n=4M/4F, tx-j n=4M/3F), and 24 weeks (C3H n=10M/12F, tx-j n=11M/11F). B: Total protein liver lysate densitometry analyses for HDAC4 and HDAC5 protein expression in 16-week old Atp7b-/-mice (n=7M/4F) and wild-type (WT, n=6M/5F). C: Immunohistochemical analysis of 24-week old C3H and tx-j mouse livers for HDAC5 (red) and DAPI (blue). Images display cytosolic and nuclear HDAC5 localization. White arrows indicate nuclei; scale bar = 50 µm. Bar graphs represent HDAC5 optical density in cytosol, nuclei, and the nucleus/cytosol ratio; n=3 mice/group, 60 cells/mouse. D: HDAC5 immunoblot of HepG2 cell lysates treated with CuSO4 (0 to 100 µM; n=3 per treatment) for 24 hours. In this figure and all subsequent figures: C3H=C3HeB/FeJ control mice with normal copper metabolism, tx-j=C3He-Atp7btx-j/J Jackson Laboratory toxic milk mouse model of Wilson disease (WD).

We then compared HDAC5 subcellular localization in liver tissue of C3H and tx-j mice by immunohistochemistry, observing reduced total HDAC5 signals in tx-j mice compared to C3H for both nuclear and cytoplasmic fractions (Figure 1C, S2C). Also, the HDAC5 signal intensity of the nucleus/cytosol ratio was significantly reduced in tx-j mice.

In an *in vitro* WD model, copper-loaded HepG2 cells also demonstrated reduced HDAC5 levels compared to control, indicating a direct effect of copper on reduced HDAC5 levels, independent from hepatic inflammation or fibrosis (Figure 1D). HDAC5-activator isoproterenol, previously shown to cause decreased hepatic and cardiac copper levels in male Wistar rats (22), increased HDAC5 levels in a dose-dependent manner when applied to copper-loaded HepG2 cells (Figure S2D). Together, these results indicate *Atp7b* mutation-induced copper accumulation reduces class IIa histone deacetylases HDAC4 and HDAC5 in WD.

### Activation of AMP-activated protein kinase (AMPK) signaling is correlated with HDAC5 phosphorylation in tx-j mice

We hypothesized the AMPK-HDAC5 regulatory pathway, previously studied in diabetes and cardiovascular disease (23), may be involved with the reduction of HDAC5 levels in WD mice, since activated AMPK signaling has been reported in animal models of copper accumulation (24). When AMPK is phosphorylated (pAMPK), it becomes active and can phosphorylate HDAC5 (pHDAC5) in the nucleus, marking HDAC5 for cytosolic exportation and possible degradation. AMPK and pAMPK protein and transcript levels were examined in livers of 24-week old tx-j mice compared to C3H (Figure 2A, 2B, S3A). AMPK and pAMPK protein and transcript levels were significantly increased in tx-j mice compared to C3H, consistent with the hypothesis. When compared to total protein levels, pAMPK and pHDAC5 ratios were significantly increased in tx-j mice relative to total AMPK and total HDAC5, respectively, whereas no significant difference was observed in C3H mice (Figure 2C).

**Figure 2.**
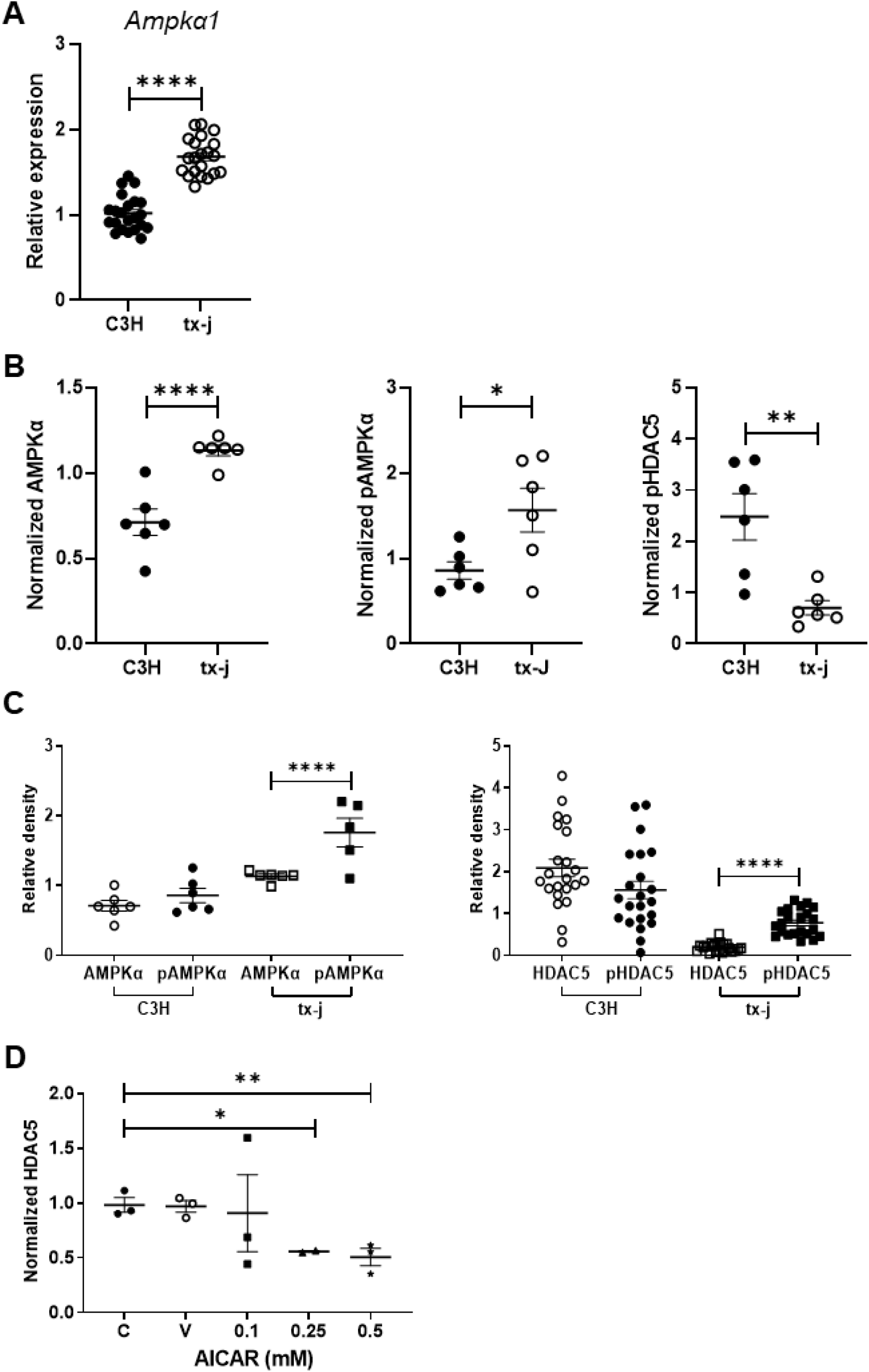
Activation of AMPKα signaling in 24-week old tx-j mice and HepG2 cells. Data are represented as means ± SEM, and asterisks indicate statistical significance determined by Student’s t-test (*p< 0.05, **p< 0.01, and ****p< 0.0001). A: Transcript levels of Ampka1 normalized to Gapdh (C3H n=10M/12F, tx-j n=11M/11F). B: Immunoblot densitometries of total AMPKα, phosphorylated AMPKα (pAMPKα), and phosphorylated HDAC5 (pHDAC5) in total protein liver lysates obtained from 24-week old C3H (n=3M/3F) and tx-j (n=3M/2F). Densitometry values were normalized to β-actin. C: Relative density comparisons of total AMPKα to pAMPKα and total HDAC5 to pHDAC5 normalized to β-actin in 24-week old C3H and tx-j mice. D: HDAC5 immunoblot densitometry analysis, normalized to β-actin, of HepG2 cell lysates treated with 50 µM CuSO4 for 24 hours followed by 5-aminoimidazole-4-carboxamide-1-β-D-ribofuranoside (AICAR) treatment (0-0.5 mM; n=3 per dose), an AMPK activator, for 24 hours. C=control, V=treated with vehicle (DMSO) only.

To determine whether activation of endogenous AMPK results in decreased HDAC5, HepG2 cells were loaded with CuSO_4_ followed by treatment with 5-aminoimidazole-4-carboxamide-1-β-D-ribofuranoside (AICAR), an AMPK activator, for 24 hours. AICAR treatment decreased HDAC5 protein levels in a dose-dependent manner (Figure 2D, S3B).

These results confirm activated AMPK is associated with increased pHDAC5 in tx-j mice compared to C3H and is the likely explanation for lower tx-j levels of HDAC5.

### HDAC4/HDAC5 deficiency in WD models is restored through copper chelation and dietary methyl donors

We next tested whether or not deficient HDAC5 levels in tx-j mice could be restored through interventions that reduce copper levels or increase methyl group donors. Twelve-week provision of PCA (copper chelator) was associated with a significant increase in hepatic HDAC4 and HDAC5 levels, similar to choline supplementation (methyl group donor). However combined treatment of PCA and choline only restored HDAC4 levels and not HDAC5 (Figure 3A).

**Figure 3.**
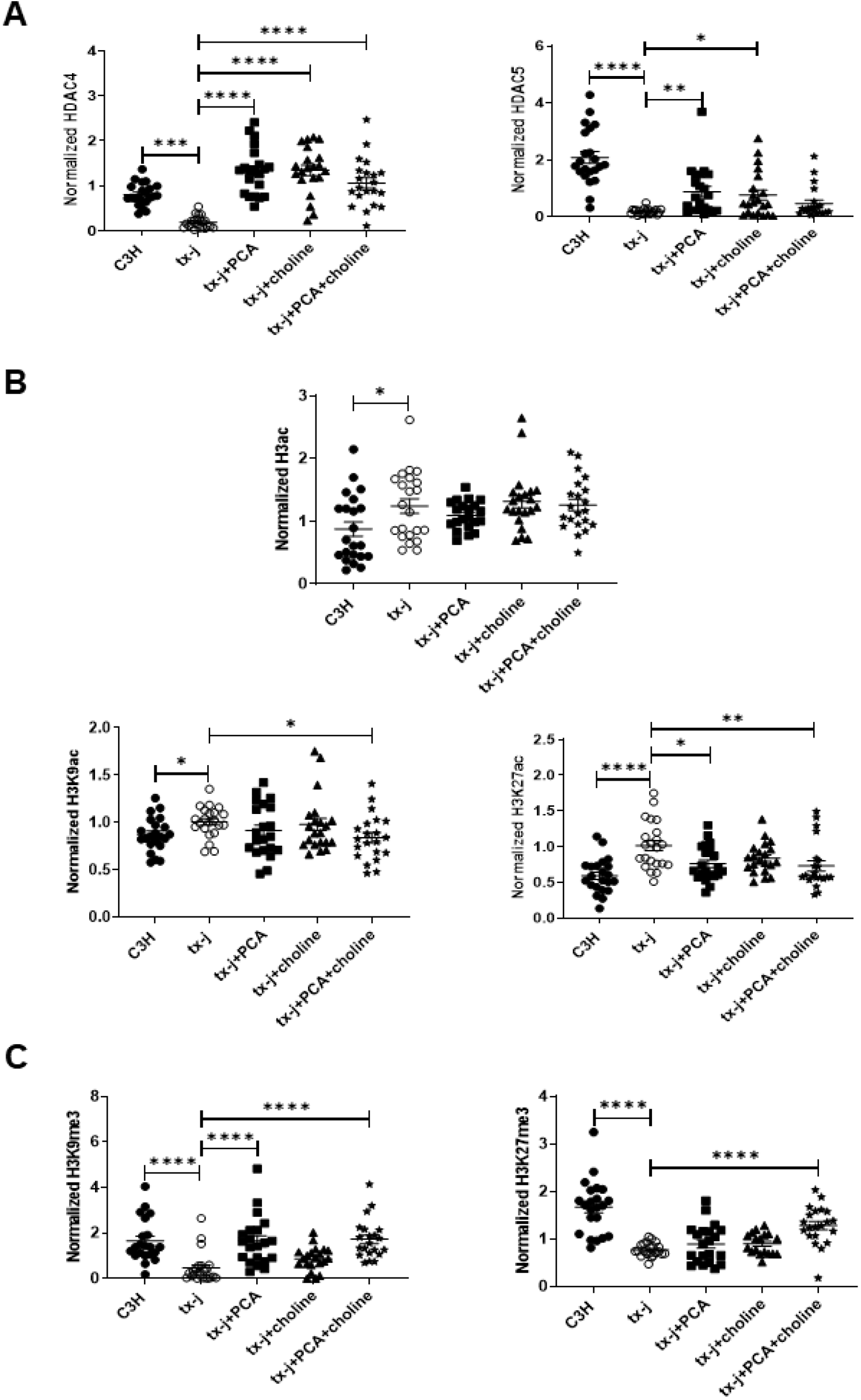
HDAC4 and HDAC5 downregulation is associated with increased histone acetylation and impaired methylation. Total protein liver lysate analyses of 24-week old C3H (n=10M/12F) and tx-j mice (n=11M/11F), and tx-j mice treated with PCA (n=11M/10F), choline (n=8M/13F), and PCA+choline (n=10M/11F). Immunoblot densitometries of HDAC4, HDAC5, H3ac, H3K9ac, H3K27ac, H3K9me3, and H3K27me3 were normalized to β-actin. Data are represented as means ± SEM and statistical analyses were performed with Kruskal-Wallis one-way ANOVA followed by uncorrected Dunn’s test (*p< 0.05, **p< 0.01, ***p< 0.001, and ****p< 0.0001).

Acetylation and methylation status in response to PCA or choline treatments were examined by assessing total histone 3 acetylation as well as histone acetylation and methylation marks H3K9 and H3K27. Consistent with reduced HDAC4 and HDAC5 levels, H3ac, H3K9ac, and H3K27ac levels were increased in livers of tx-j mice compared to C3H, with H3K27ac changes the most prominent. PCA intervention counteracted histone acetylation and methylation mark effects, with restoration of H3K9ac and H3K27ac to C3H levels most apparent with combined treatments; however, total H3 acetylation remained unaffected (Figure 3B, S4A). This implies the H3 acetylation response to PCA is specific to H3K9 and H3K27. H3K27me3 and H3K9me3 were significantly reduced in tx-j compared to C3H mouse livers, and PCA restored H3K9me3 and H3K27me3 to levels observed in C3H livers (Figure 3C). In contrast, choline supplementation alone was not associated with any changes in histone acetylation and methylation marks. The reduced levels of H3K9ac and H3K27ac and increased levels of H3K9me3 and H3K27me3 in the PCA+choline group compared to untreated tx-j, then, were likely an effect of PCA alone and not the combined effect of the two interventions. As histone acetylation is dynamically regulated by the balance between histone acetyltransferase (HAT1) and HDAC activities, it is worth noting *Hat1* hepatic transcript levels were increased in tx-j mice at baseline with no significant effects induced by the interventions (Figure S4B).

### Chromatin immunoprecipitation sequencing (ChIP-seq) for H3K9ac and H3K27ac integrated with RNA-seq reveal altered epigenetic regulation and transcript levels of genes involved in energy metabolism and liver regeneration

To identify genes regulated by the epigenetic alterations to histone acetylation in tx-j mice, we conducted and integrated analyses of H3K9ac and H3K27ac by ChIP-seq and gene expression by RNA-seq in liver. Differential H3K27ac peaks in tx-j compared to C3H liver were mapped to more than 8,000 genes (I) (Table S1), but the number of significant differential H3K27ac-associated genes was reduced when tx-j mice were compared to those treated with PCA (II), choline (III), or the combination of the two interventions (IV) (Figure 4A). Differential H3K9ac peaks between tx-j and C3H liver were mapped to >500 genes (Table S5). Similar to the H3K27ac analysis, transcriptome analysis revealed more than 10,000 genes differentially expressed in tx-j compared to C3H liver (I) (Table S6), with fewer differentially expressed genes comparing groups of tx-j mice in response to interventions (II, III, IV) (Figure 4B). When comparing untreated tx-j to PCA-treated tx-j liver, 40 genes were found to be differentially enriched for H3K27ac, whereas 2,554 genes were differentially expressed (Table S2 and S7). Increased methyl group availability through choline supplementation showed a small effect on histone acetylation-mediated gene transcription with 112 genes differentially acetylated and 6 genes differentially expressed when comparing untreated to choline-treated tx-j mice (Table S3 and S8). In PCA+choline-treated mice vs. untreated, only 30 H3K27ac genes were enriched, whereas 2,776 genes were differentially expressed (Table S4 and S9). As shown in volcano plots, there were significant differences between tx-j and C3H control mice for both H3K27ac ChIP-seq and RNA-seq, often in the same direction as either co-upregulated or co-downregulated genes (Figure S5).

**Figure 4.**
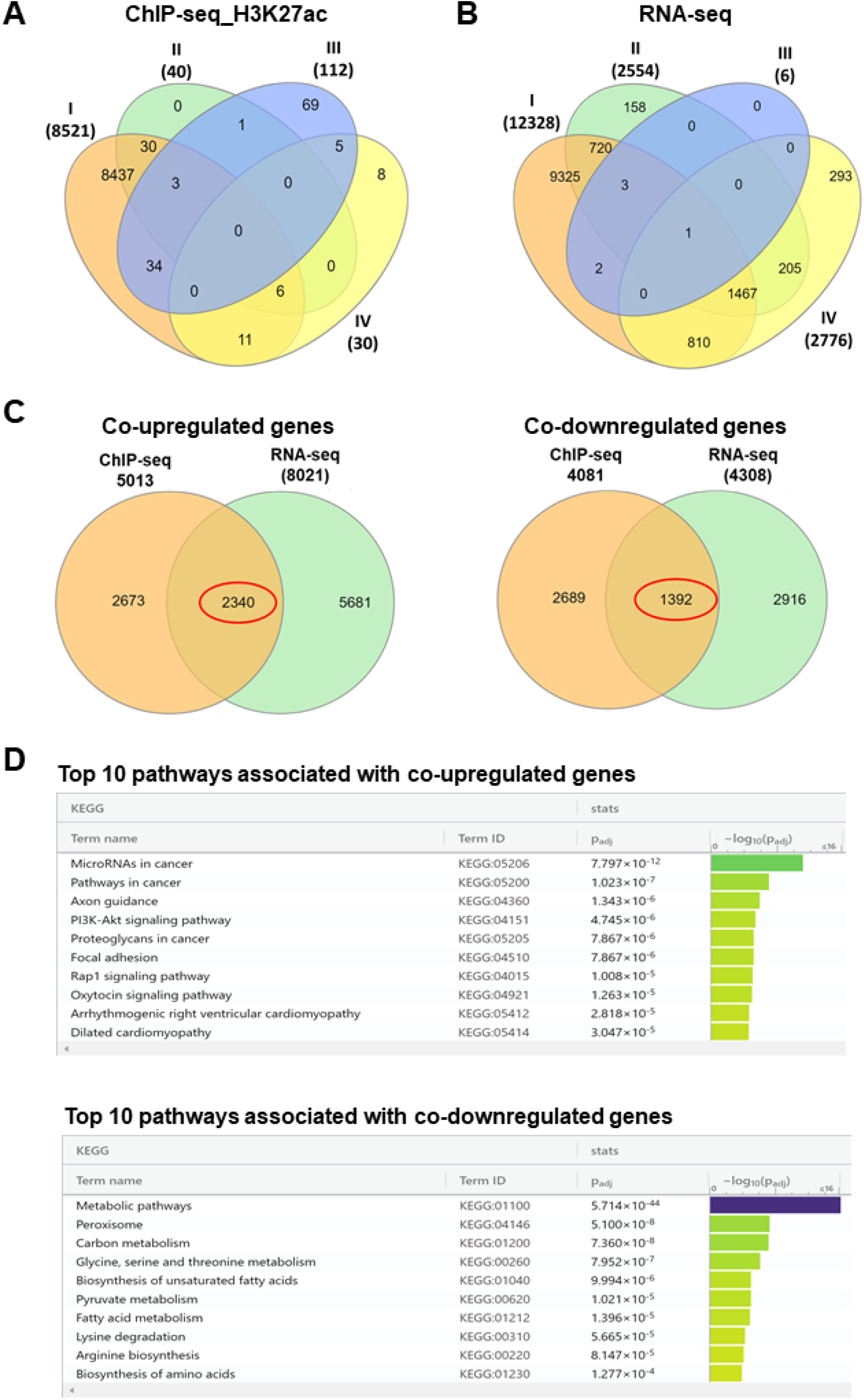
H3K27ac-associated ChIP-seq and RNA-seq data analyses in the liver of tx-j mice. ChIP-seq and RNA-seq was performed in livers of 24-week old C3H, tx-j, tx-j+PCA, tx-j+choline, and tx-j+PCA+choline groups (n=3M/3F per group). A and B: Venn diagrams showing numbers of H3K27ac-associated differentially enriched genes detected by ChIP-seq and differentially expressed genes by RNA-seq. I = C3H vs. tx-j, II = tx-j vs. tx-j+PCA, III = tx-j vs. tx-j+choline, IV = tx-j vs. tx-j+PCA+choline. C: Numbers of common significantly co-upregulated and co-downregulated genes between ChIP- and RNA-seq analyses. D: Pathway enrichment analysis showing the top 10 significantly associated pathways with differentially expressed co-upregulated and co-downregulated genes between ChIP- and RNA-seq by g:Profiler.

When ChIP-seq and RNA-seq analyses were overlapped, 2,340 genes were co-upregulated and 1,392 were co-downregulated between C3H and tx-j liver (Figure 4C, Table S10). Pathway analysis of co-upregulated and co-downregulated genes from ChIP-seq and RNA-seq analyses using g:Profiler (25) identified target genes related to the PI3K-AKT signaling pathway (55 genes), including *Fgfr1, Bcl2l1, Ccnd2, Col4, Myb*, and *Lama5*; one-carbon metabolism (29 genes), including *Gck, Aldh6a1, Cat, Eno, Adh5, Acat2*, and *Sdh*; fatty acid metabolism (18 genes), including *Scd1, Acaa2, Fasn, Scp2*, and *Acat2*; lysine degradation (16 genes), including *Kmt5a, Gcdh, Nsd1, Kmt2b, Setmar, Ehmt2, Acat2*, and *Setd1a*; and biosynthesis of amino acids (20 genes), including *Shmt2, Acy1, Eno1, Mat1a*, and *Pklr* (Figure 4D, Table S11).

Next, we specifically examined differential expression of 1,623 genes encoding transcription factors (TF) in tx-j compared to C3H mice. As shown in the Venn diagram, 174 TF genes were shared between tx-j co-upregulated genes and 80 TF genes were shared between tx-j co-downregulated genes when compared against total known TF genes in mice (Figure 5A, Table S12). ChIP-seq peaks revealed notably different enrichment of H3K27 acetylation sites for several upregulated and downregulated TF genes in tx-j mice compared to C3H (Figure 5B). Differentially expressed TF genes were also investigated for functional relevance by LISA (epigenetic Landscape In Silico deletion Analysis) (Figure S6). Top enrichments included transcription factors known to play a role in liver function/development (FOXA2, HNF4A, RXRA) and liver lipid metabolism (PPARα).

**Figure 5.**
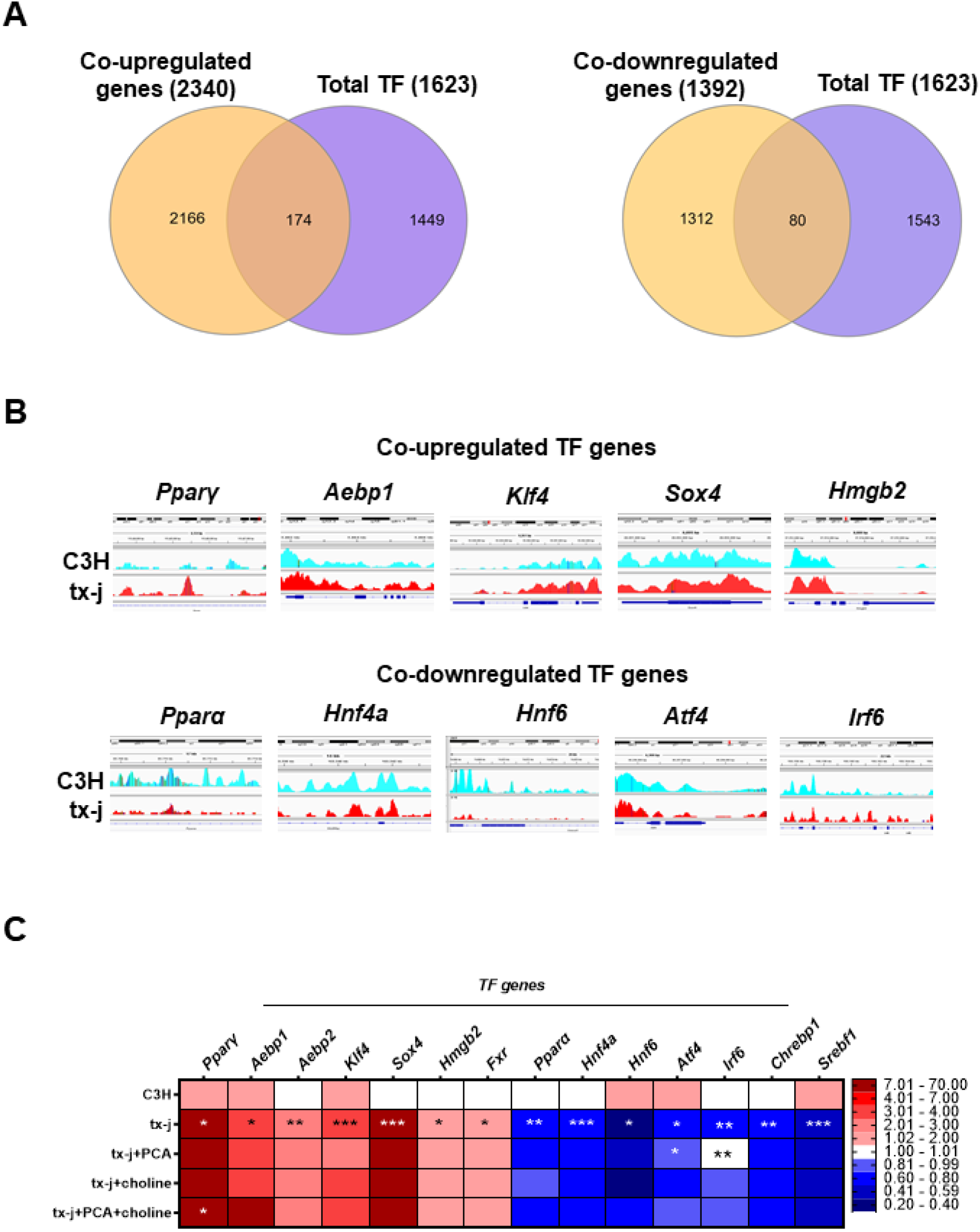
Transcription factors within differentially expressed genes between C3H and tx-j mice. Using the ChIP-seq and RNA-seq data from livers of 24-week old C3H and tx-j mice (3M/3F per group), differentially expressed genes in ChIP- and RNA-seq analyses were overlapped with total known mouse transcription factors. A: Venn diagram showing numbers of common co-upregulated (174) and co-downregulated (80) transcription factor genes. B: ChIP-seq peaks showing enrichment of H3K27ac for transcription factors, selected by their involvement in liver and metabolic diseases. C: Heatmap displaying qPCR gene transcript levels of representative mouse transcription factors. Relative expression data are normalized to Gapdh (3M/3F per group). Data are represented as means ± SEM and statistical analyses were performed with Kruskal-Wallis one-way ANOVA followed by uncorrected Dunn’s test (*p< 0.05, **p< 0.01, and ****p< 0.0001) for C3H vs. tx-j and tx-j vs. tx-j+PCA, choline, or combined treatments

To gain mechanistic insight into histone deacetylation and acetylation-/methylation-associated TF regulation, we examined the transcript levels of key TF genes involved in liver development (*Hnf4a, Hnf6*), liver fibrosis (*Aebp1* and *2, Smad2, Hmgb2*), steatosis and lipogenesis (*Pparγ, Sox4, Fxr, Chrebp, Pparα, Irf6, Srebf1*), oxidative stress (*Klf4, Atf4*), and apoptosis (*Irf6*). Gene expression analysis revealed significantly upregulated TF genes, including, *Pparγ, Aebp1* and *Aebp2, Sox4, Klf4, Hmgb2*, and *Fxr* while significantly downregulated genes included *Pparα, Hnf4a, Hnf6, Irf6, Atf4, Chrebp*, and *Srebf1* in tx-j compared to C3H mice (Figure 5C). Copper chelation with PCA significantly increased transcript levels of *Irf6* and *Atf4* while choline supplementation did not show any significant effect. PCA+choline only elicited a significant response from *Pparγ*.

### HDAC5 levels in tx-j mice are modulated by high fat dietary intervention

HDAC5 is known to be affected by nutrient availability and the metabolic state (26). Given this trait and the associated lipid-related metabolic pathways from the acetylome/transcriptome analyses, tx-j and C3H mice were challenged with a 60% kcal fat diet (HFD) for 8 days and livers collected at 9 weeks of age. The 9-week time point was chosen as tx-j mice do not yet have significant liver disease at this early stage, therefore, changes seen in HDAC levels would be independent of liver damage. Tx-j mice on control chow demonstrated significantly lower levels of triglycerides compared to C3H (Figure 6A). Both tx-j and C3H mice on HFD had elevated hepatic triglycerides and cholesterol compared to chow, confirming the HFD had a rapid and detectable effect in the liver. Following the same pattern, HDAC5 protein levels trended lower in tx-j mice on control chow compared to C3H; however, after 8 days of HFD, tx-j HDAC5 protein levels were significantly increased (Figure 6B, S7A).

**Figure 6.**
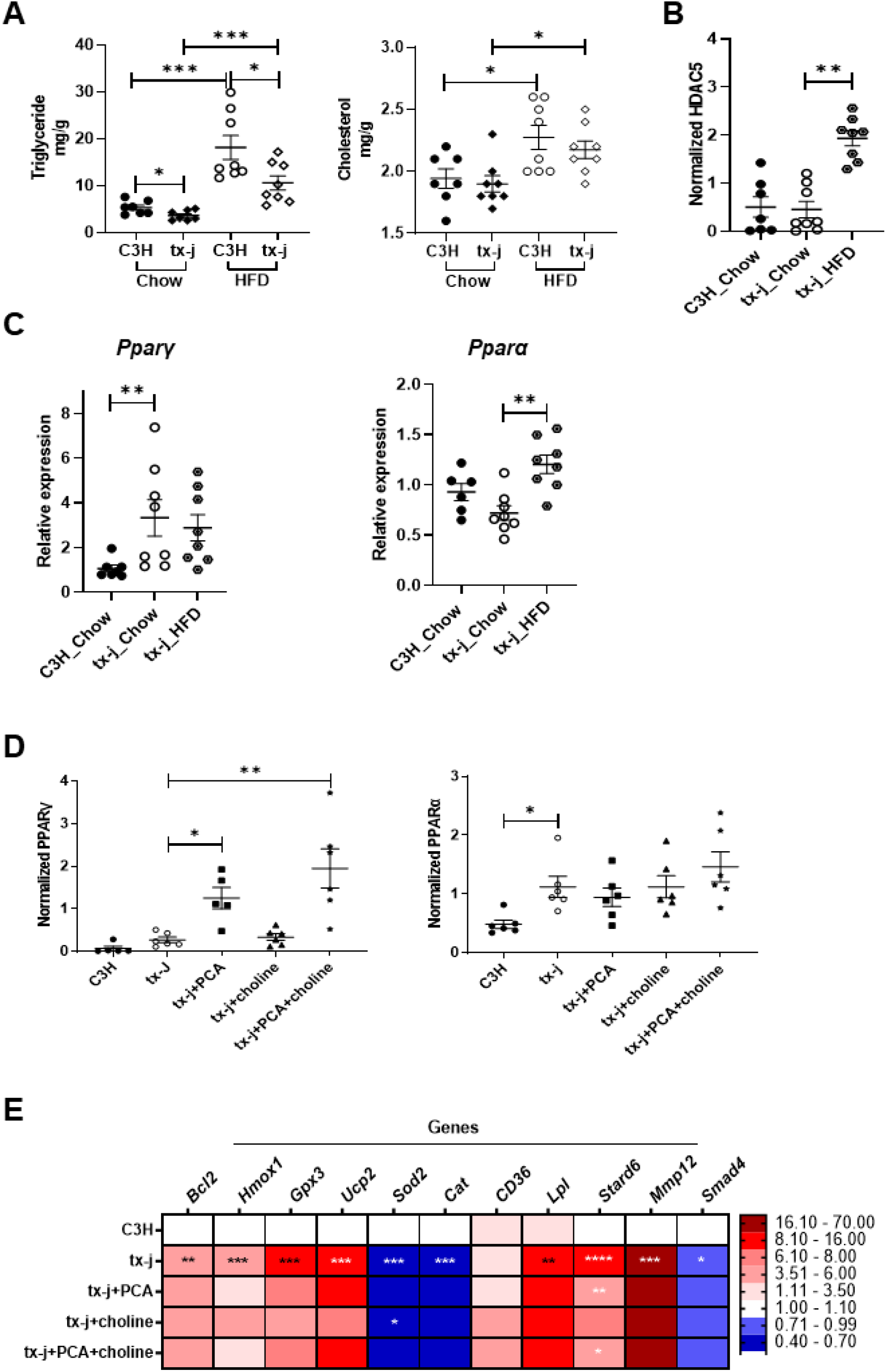
Dietary modulation (high fat, choline, PCA) of HDAC5 and metabolic regulators PPARα and PPARγ. Data for A-D are represented as means ± SEM. Statistical significance for all analyses was determined by Student’s t-test for comparison between two groups and Kruskal-Wallis one-way ANOVA followed by uncorrected Dunn’s test for multiple groups (*p< 0.05, **p< 0.001, ***p< 0.001, and ****p< 0.001). A: Liver triglyceride and total cholesterol of mice on chow (C3H n=4M/3F, tx-j n=4M/4F) compared to C3H and tx-j (n=4M/4F each) fed a 60% kcal fat diet (HFD). Mice were challenged with HFD for 8 days and tissues collected at 9 weeks of age. B: HDAC5 protein expression in total protein liver lysate from mice on chow or HFD. Immunoblot densitometry analysis was normalized to β-actin. C: Liver transcript levels of Pparγ and Pparα normalized to Gapdh in mice on chow or HFD. D. Immunoblot densitometries of PPARγ and PPARα, normalized to β-actin, in total protein liver lysate of 24-week old C3H (n=3M/3F), tx-j (n=3M/2F), tx-j+PCA (n=3M/3F), tx-j+choline (n=3M/3F), and tx-j+PCA+choline (n=3M/3F). E: Heatmap representing qPCR transcript levels of PPARα- and PPARγ-related genes measured in livers of 24-week old C3H vs. tx-j and tx-j vs. tx-j+PCA, choline, or combined treatments (n=3M/3F per group). Relative expression data are normalized to Gapdh.

Peroxisome proliferator-activated receptor alpha (PPARα) and gamma (PPARγ) were selected from the TF gene analysis for further examination as they are key regulators of lipid metabolism. PPARα and PPARγ were also shown to be associated with steatosis and the antioxidant response, and exhibited decreased and increased protein levels, respectively, concomitant with liver damage severity in patients with WD (27). During an early stage of disease progression (9 weeks of age), *Pparγ* transcript levels were significantly higher in tx-j than C3H mice and *Pparα* transcript levels were reduced, but 8 days of HFD were associated with significantly increased *Pparα* expression (Figure 6C). At a later stage of disease progression (24 weeks of age) in tx-j mice, PPARα and PPARγ (non-significant) protein levels were increased in tx-j compared to C3H mice (Figure 6D, S7B). In response to PCA, PPARγ levels were further increased compared to untreated tx-j, whereas PPARα was unaffected by PCA and/or choline. Moreover, transcript levels of PPARα- and PPARγ -associated genes *Bcl2, Hmox1, Gpx3, Ucp2, Lpl, Stard6*, and *Mmp12* were significantly increased while *Sod2, Cat*, and *Smad4* were decreased in tx-j mice compared to C3H. These findings indicate increases in fibrosis (*Smad4*), anti-apoptosis (*Bcl2, Mmp12*), antioxidant responses (*Hmox1, Ucp2, Gpx3, Cat, Cd36*), uptake of fatty acids and beta oxidation (*Cat, Cd36, Lpl*), and lipogenesis (*Stard6, Sod2*) in tx-j mice. PCA+choline treatment had a limited effect on gene expression, except on *Stard6*. Of note, *Hmox1* was significantly increased in tx-j mice after 8 days of HFD (Figure S7C). Overall, these data suggest involvement of HDAC5 in the regulation of PPARα and PPARγ and their associated pathways in WD, and its potential to be modulated by dietary lipid intake.

## Discussion

The clinical, metabolic, and liver pathology features of WD indicate this condition, although rare, presents characteristics of more common liver diseases, including nonalcoholic fatty liver disease. Even though copper accumulation is clearly the primary underlying mechanism of liver damage in WD, the intersection of dietary, epigenetic, bioenergetic, and metabolic defects are contributing, amplifying, and potentially acting as priming factors in the development of hepatic and extrahepatic manifestations. Therefore, it is of crucial importance to identify the modulating elements connecting genetic, epigenetic, and metabolic pathways in WD, and we propose HDACs as major players in this role. The main findings of our study are, first, the identification of decreased HDAC4 and HDAC5 levels in livers of tx-j mice starting at early stages of hepatic copper accumulation and progressing with age-related worsening of liver disease. Second, reduced levels of HDACs are associated with dysregulation of histone acetylation marks H3K9ac and H3K27ac, corresponding with effects on gene expression. Furthermore, HDAC and histone acetylation alterations were counteracted by dietary supplementation of fat or methyl groups, or copper chelation by PCA. Finally, the genes affected by H3K27 acetylation in WD are involved in metabolism regulation. In recent years, expanding data have shown HDACs are key regulators of glucose homeostasis, and cardiovascular and endothelial function (28, 29). HDACs are also known to be regulated by dietary factors as well as fasting versus feeding status. In agreement with available research, we demonstrated class II HDAC levels are regulated by multiple factors including copper levels, availability of methyl groups, and dietary fat.

Several studies have shown class IIa HDACs are regulated by AMPK via phosphorylation (30). Class IIa HDACs are highly regulated by their phosphorylation/de-phosphorylation status with consequent modulation of gene transcription. HDAC5 molecules are phosphorylated at a number of serine residues which provide binding sites for chaperone proteins to export HDAC from the nucleus to cytosol (31). Sequence analysis indicated phosphorylation sites are largely preserved between HDAC isoforms. In mouse livers, class IIa HDACs are phosphorylated by AMPK family kinases (32). Hepatocyte-specific knockdown of class IIa HDACs via RNAi-induced deacetylation of FOXO led to inhibition of FOXO target genes with consequent lower blood glucose levels and increased glycogen storage. Salt-inducible kinase-dependent de-phosphorylation of HDAC4 increases lipolysis through FOXO1 inactivation that eventually leads to depletion of accumulated triglyceride. Conversely, HDAC4 downregulation restores lipid accumulation (33, 34). In mouse primary hepatocytes, HDAC5 plays a role in integrating the fasting state and ER stress signals to regulate hepatic fatty acid oxidation. This suggests HDAC5 could be a target for the treatment of obesity-associated hepatic steatosis (20). Taken together, these studies indicate central roles for class II HDACs in metabolic processes, including glucose homeostasis, energy metabolism, and lipogenesis.

While the primary downstream effects of class II HDACs impact glucose and lipid metabolism, they also have detrimental systemic effects. A mouse model with a liver-specific knockdown of class IIa HDACs, including HDAC4, HDAC5, and HDAC7, exhibits not only carbohydrate and lipid alterations in hepatocytes, but also systemic manifestations involving kidney and spleen abnormalities (35). This suggests the combination of reduced class IIa HDAC levels and increased copper levels could explain WD systemic manifestations which span from brain to heart and kidney involvement. A key element in interpreting our data is the time-course study on HDAC5 levels. Reduced hepatic HDAC5 levels were observed starting at day 6 post-partum, when hepatic copper levels are normal. The progression of hepatic copper accumulation and hepatic inflammation and fibrosis accompanied progressive reductions in tx-j HDAC levels, though it is evident reduced HDAC levels are intrinsic to the tx-j mouse model and likely the result of impairments in development and regeneration mechanisms (10). These data are therefore more consistent with the reported proteasome-dependent degradation of phosphorylated class IIa HDACs in the cytosol (36) than cytosolic protection by 14-3-3 binding (37).

The regulation of HDAC5 and HDAC4 is complex. We propose the simplest mechanism explaining reduced HDAC5 in WD and its subsequent impact on transcription is phosphorylation by AMPK, leading to relocation from the nucleus to the cytosol and followed by degradation. Acute oxidative stress, ATP depletion, and transiently elevated calcium are conditions thought to initiate phosphorylation of class IIa HDACs (38). Notably, AMPK is a kinase well-known to be activated to various degrees by all these stimuli (39). The phosphorylation of threonine-172 activates AMPK and is associated with increased transcript levels of genes related to metabolism. As shown in our study and reported by previous *in vitro* studies, AMPK can phosphorylate HDAC5 (40). Our current data showed increased ratios of phosphorylated to unphosphorylated AMPK and HDAC5 in WD mice (Figure 2). Moreover, hepatocyte treatment with an AMPK activator, AICAR, was associated with reduced expression of HDAC5, likely due to a nucleus-to-cytosol pHDAC5 shift.

Our results reveal HDAC5 levels are affected by dietary factors such as availability of methyl groups via choline supplementation and excess dietary fat. This likely has an effect on the risk of developing hepatic steatosis as well as systemic metabolic manifestations. Reduced class IIa HDAC4 and HDAC5 might play a central role in the regulation of H3K9 and H3K27 acetylation/methylation. Of note, HDAC5 knockdown was shown to increase H3K9 acetylation in mouse C3H10T1/2 cells (41), similar to tx-j vs. C3H mice in our study. Altered HDAC4/5 regulation coincides with copper overload as shown in tx-j mice where acetylation and methylation are restored to control levels in response to copper chelation (Figure 3). H3K9ac and H3K27ac are associated with transcriptionally active chromatin, whereas H3K9me3 and H3K27me3 are associated with an inactive heterochromatin state. In our studies, this histone acetylation-methylation shift may lead to transcriptional reprogramming with consequent aberrant expression of numerous genes in WD.

ChIP-seq for H3K27 acetylation and RNA-seq analyses confirmed the regulation of hepatic metabolic pathways through histone acetylation mechanisms. In particular, the PI3K-AKT pathway was co-upregulated in tx-j compared to CH3 liver across both analyses. This pathway is of interest as it is extensively shown to be upregulated during the progression of NAFLD with associated risk of liver cancer (42, 43). Other co-regulated pathways included one-carbon metabolism, which plays an important role in regulating energy metabolism and immune function in fatty liver disease and fibrosis (44). Our previous study also revealed copper-mediated inhibition of SAH hydrolase and the consequent accumulation of SAH leads to global DNA hypomethylation in tx-j mice (12). Our current findings corroborate and add to previous reports indicating an acetylation/methylation shift is involved in the regulation of metabolic pathways and related TF genes, including *Pparα, Pparγ, Chrebp, Srebf1, Fxr, Lxr, Rxra*, and *Stard6* (Figure 5C, Table S13). Some important differentially regulated genes involve the following pathways: cell regeneration (*Hnf4a, Hnf6*, and *Myb*); cell cycling, development, and differentiation (*Fgfr1, Bcl2, Sox, Timp2, Gata2*, and *Klf4*); fatty acid metabolism (*Scd1, Fads, Fasn*, and *Acat2*); and oxidative and ER stress (*Hmox1* and *Atf4*). Our previous studies on DNA methylation in liver from WD patients showed hypermethylated WD liver differentially methylated regions were enriched in liver-specific enhancers, flanking active liver promoters and binding sites of liver developmental transcription factors, including HNF4A, RXRA, and FOXA2 (9), which were recapitulated in the top 10 transcription factors enriched in this RNA-seq dataset. This suggests convergence of DNA methylation and transcription factor regulation of gene expression in WD. Two of the most interesting targets are PPAR isoforms PPARα and PPARγ, known to regulate lipid and glucose homeostasis through their coordinated activities in liver, muscle, and adipose tissue (45). Our findings of increased PPARα and PPARγ in tx-j mice correspond to a study indicating changes in PPARα and PPARγ in WD patients (27). Previous studies have shown PPARα increases with mild liver damage and decreases with moderate or greater liver damage, while PPARγ was increased in WD patients concomitantly with the progression of liver damage. No previous studies have focused on PPAR signaling in WD. Many studies report the anti-inflammatory, antioxidant, and anti-apoptotic roles of PPARα and γ (46-49), which are further supported by our data of significantly increased PPARγ gene and protein expression after PCA treatment.

Activation of PPARα and PPARγ signaling is known for decreasing triglyceride and cholesterol levels in liver and plasma of mice and humans (50). Corroborating these effects, we also found liver (Figure 6A) and plasma (data not shown) triglyceride and total cholesterol levels reduced in tx-j mice on chow as well as HFD compared to control. Furthermore, the observed PPARγ effect on antioxidant gene expression of *Hmox1, Ucp2, Gpx3*, and *Cd36*, and anti-apoptotic gene *Bcl2*, could be part of a protective mechanism against copper-mediated oxidative stress in WD. A previous study showed mice fed a high fat diet for 4 weeks exhibited increased HDAC5 levels in medial hypothalamus (51); we confirmed that even a short-term, 8-day high fat challenge induced a dramatic HDAC5 response. Interestingly *Hmox1* (Figure S7C) and *Pparα* (Figure 6C) showed significant upregulation after short-term HFD exposure in tx-j mice. Bringing together all of the studied elements, a working model of HDAC5 regulation and impact on histone acetylation and subsequent gene regulation is proposed in Figure 7.

**Figure 7.**
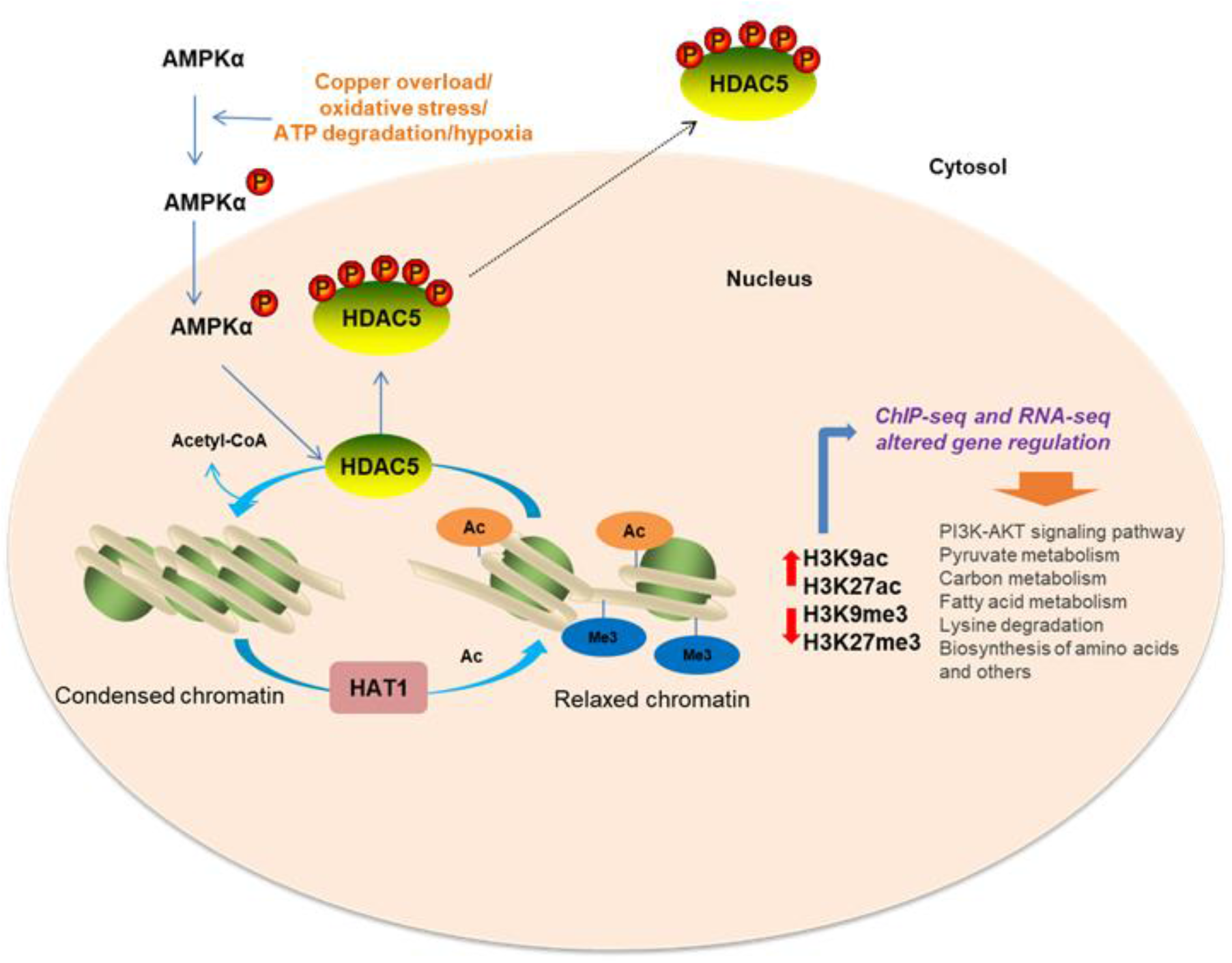
Proposed schematic of HDAC5-mediated H3K9ac and H3K27ac regulation of gene expression and affected biological pathways in WD. In WD, copper overload and oxidative stress might lead to phosphorylation of AMPK (active form). Increased phosphorylated AMPK could then phosphorylate HDAC5 which is subsequently exported to the cytosol. Lack of nuclear HDAC5 and increased histone acetyltransferase (HAT1) might cause an increase in acetylated histones (H3K9ac and H3K27ac) and decrease methylated histones (H3K9me3 and H3K27me3) with subsequent altered regulation of genes in WD. Our results show HDAC5 impacts the acetylation/methylation balance and serves as a critical regulator of genes central in metabolic regulation. ChIP-seq and RNA-seq revealed 3,732 differentially expressed genes in the tx-j mouse model of WD and the enrichment analysis of these genes included pathways related to lysine degradation, fatty acid metabolism, carbon metabolism, pyruvate metabolism, and signal transduction.

One disappointing but still insightful finding of our study is the limited effect of PCA, choline, and combined treatment on the number of genes regulated by histone acetylation. This could be interpreted as a potential explanation for the lack of complete response to anti-copper treatment often observed in patients with WD. These findings may be due to the relatively short duration of intervention; copper chelation alone is not enough to reverse metabolic perturbations; and dietary choline can be shunted to other pathways in addition to DNA methylation. It remains unknown if metabolic aberrations in WD are independent or a direct result of copper accumulation. It is clear, however, that removal of excess copper alone cannot restore all epigenetically compromised pathways in WD. More interventions targeting HDAC5 and other identified dysregulated pathways are warranted and could serve as adjunct treatments. An appealing facet of HDAC5 is its ability to be modulated rapidly through diet, making it a potential target particularly for early intervention to prevent severe liver damage and associated metabolic complications. Among the identified TFs, PPARγ could emerge as a therapeutic target for fatty liver in WD, especially during the course of treatment with PCA or other copper chelators. Ultimately, class IIa HDAC4 and HDAC5, as key metabolic elements, could represent a regulatory interface between genetic and metabolic factors affecting the varied phenotypic presentations and liver disease manifestations in WD.

## Materials and Methods

### Animals

C3HeB/FeJ (C3H) control, Jackson Laboratory toxic milk model C3He-Atp7btx-J/J (tx-j), and *Atp7b*^-/-^colonies were maintained at 20–23°C, 40–65% relative humidity, and a light cycle of 14 h light/10 h dark. Mice were maintained ad libitum on deionized water and LabDiet chow (Purina Mills, Inc.). C3H and tx-j mice were each maintained as homozygous colonies. *Atp7b*^-/-^mice were maintained by a heterozygous breeding colony which generated both homozygous mutant and wild-type controls. Tx-j mouse milk lacks sufficient copper for neonatal growth and development beyond day 10 on average hence all tx-j pups were fostered to a lactating C3H dam by day 7 post-partum with the exception of PPD6 pups from which tissues were collected prior to fostering. Progeny were weaned between 21-28 days of age. A subgroup of the 9-week tx-j cohort was challenged with a 60% kcal fat diet (Research Diets), 8 days prior to tissue collection. At PPD6, 9, 12, 16, and 24 (C3H only) weeks of age, mice were euthanized and livers were flash-frozen in liquid nitrogen then stored at -80°C until further analysis. All tissues were collected in the morning, between 3-6 hours after lights went on in the animal room. All groups had relatively equal male/female representation.

For the 24-week tx-j cohort, tx-j females were fed either control Teklad 2020 chow (Envigo) or choline-supplemented Teklad 2020 chow (5 g/kg diet) 2 weeks prior to mating. To circumvent the homozygous recessive mutation in tx-j dams which causes copper-deficient milk production, heterozygous tx-j breeders were used for this cohort to generate homozygous mutant tx-j progeny. Three days prior to mating, male breeders were fed whichever diet matched their prospective female to acclimate. Dams continued with their respective diets through mating, gestation, and lactation. DNA samples were obtained from pups between 8-12 days post-partum for genotyping. Genotyping was performed by the UC Davis Mouse Biology Program via Taqman allelic discrimination. Progeny were weaned between 21-27 days of age onto the same diet as their dam. From 12 to 24 weeks of age, D-penicillamine (PCA, Sigma-Aldrich) was administered orally in the drinking water (100 mg/kg body weight/day) to a subgroup of tx-j control and choline-supplemented mice. Mice were euthanized at 24 weeks of age and livers were flash-frozen in liquid nitrogen then stored at -80°C until further analysis. All tissues were collected in the morning, between 3-6 hours after lights went on in the animal room. All groups had relatively equal male/female representation.

All mouse protocols were reviewed and approved annually by the UC Davis Institutional Animal Care and Use Committee and followed the guidelines of the American Association for Accreditation of Laboratory Animal Care. Humane care was provided according to criteria outlined in the “Guide for the Care and Use of Laboratory Animals” prepared by the National Academy of Sciences and published by the National Institutes of Health (NIH publication 86–23 revised 1985).

### HepG2 cell culture experiments

HepG2 cells were obtained from ATCC and cultured in Eagle’s minimum essential medium with L-glutamate (Quality Biological), 10% fetal bovine serum (Fisher Scientific), and 1% penicillin/streptomycin (Gibco); cells were incubated at 37°C and 5% CO2. An equal number of cells were seeded in each well of 6-well or 10-cm collagen-treated cell culture plates per treatment group and incubated overnight to attach, with subsequent media changes twice per week. Upon 70-75% confluency, cells were washed twice with PBS and treated with 100 mM CuSO4 for 24 hours. Untreated HepG2 control cells were washed twice with PBS and provided with fresh medium. The following day, copper-treated cells were washed twice with PBS and subjected to treatment with PCA (1mM), 5-aminoimidazole-4-carboxamide-1-β-D-ribofuranoside in DMSO (0.1-1mM, AICAR, Fisher Scientific), or isoproterenol (5-25µM, Sigma-Aldrich) for 24 hours.

### RNA isolation

Total RNA was extracted from frozen mouse liver (20-25 mg) and HepG2 cells (5-6 million) using the AllPrep DNA/RNA Mini Kit (QIAGEN) according to manufacturer’s instructions. Sample purity and concentration were measured by a Nanodrop spectrophotometer (Thermo Fisher Scientific); RNA integrity was evaluated by agarose gel electrophoresis. Total RNA was stored at -80°C until further use.

### Quantitative PCR

Mouse liver total RNA (5 µg) and HepG2 total RNA (4 µg) was input to reverse transcribe with the SuperScript III First-Strand cDNA Synthesis kit (Invitrogen) according to manufacturer’s instructions. Primers for mouse cDNA sequences were designed using the free online application Primer 3 (http://bioinfo.ut.ee/primer3-0.4.0/) and blasted against the mouse genome using NCBI Nucleotide BLAST to check primer specificity (http://blast.ncbi.nlm.nih.gov/Blast.cgi). The amplification efficiency (E) of all assays was calculated from the slope of a standard curve generated via 10-fold serial dilution of pooled control cDNA using the formula E = 10(−1/slope) -1. In-software melt curve analysis and agarose gel electrophoresis of the PCR product were both checked to confirm amplicon specificity. All samples (no-template control, mouse cDNA 1/25 dilution, inter-plate QC) were run in triplicate with SYBR Green Master Mix (Applied Biosystems) on a ViiA 7 Real-time PCR System (Applied Biosystems). Primer sequences used are listed in Table S13.

### Western Blot

Mouse liver samples (35-40 mg) were homogenized on ice with a hand-held OMNI TH homogenizer (OMNI International) with ice-cold RIPA lysis buffer containing Complete Mini Protease Inhibitor Cocktail and PhosSTOP Phosphatase Inhibitor Cocktail (Roche Diagnostics). In cell culture experiments, cells were lysed by adding ice-cold RIPA buffer directly to each well. Cell protein lysates were homogenized by sonication for 2-5 seconds. Both cell and tissue lysates were centrifuged for 20 minutes at 10,000 rpm and 4°C and the supernatant collected. Protein concentration was determined with the Pierce BCA Protein Assay Kit (Thermo Fisher Scientific) according to the manufacturer’s protocol. All samples and standards were plated in duplicate and read on a Synergy H1 microplate reader (Bio Tek). Equal amounts of protein (25 µg) were denatured 1:1 in 2X Laemmli buffer (BIO-RAD) at 95°C for 10 minutes then held on ice. Proteins were separated by SDS-PAGE and transferred onto a pre-activated PVDF membrane using the Trans-Blot Turbo transfer system (BIO-RAD). Membranes were blocked with 5% non-fat milk in tris-buffered saline with Tween-20 (TBST) for 2 hours then probed with the desired primary antibodies overnight at 4°C. Membranes were washed in TBST then incubated with the appropriate horseradish peroxidase-conjugated anti-rabbit or anti-mouse IgG secondary antibody for 1 hour at room temperature. Membranes were washed in TBST again and blots were developed with Clarity Western ECL Substrate (BIO-RAD). Densitometry analyses were quantified by using Fujifilm Multi Gauge software (Fujifilm USA) and normalized to GAPDH. Immunoblot images are shown in Figures S1, S2, S3, S4, and S7; antibodies and dilutions are listed in Table S14.

### Immunohistochemistry

Immunohistochemical analysis was performed as previously described (52, 53). Briefly, mouse liver tissues were fixed in paraformaldehyde, embedded in paraffin, and sectioned. Tissue sections were stained with anti-HDAC5 antibody (Santa Cruz Biotechnology) and cell nuclei were detected by 4′,6-diamidino-2-phenylindole (DAPI). Parallel staining with non-specific IgG was performed to detect background noise. Imaging was performed using an All-in-One Fluorescence Microscope BZ-X800 (Keyence); optical density analysis was performed using ImageJ (NIH). Tissues from 3 mice/group, 60 cells/mouse were analyzed.

### Liver triglycerides and cholesterol

All samples were assayed by the UC Davis Mouse Metabolic Phenotyping Center. Briefly, 100-110 mg liver was homogenized using sodium sulfate and a pestle and mortar. Chloroform and methanol were added at a 2:1 ratio and samples were incubated overnight at 4°C. Sodium chloride (0.7% solution) was added the next day and samples were incubated overnight at 4°C to separate the chloroform layer. The next day, the supernatant was aspirated, and a sample was taken from the chloroform layer and evaporated using N2 gas. The sample was reconstituted with isopropanol and reagents from Fisher Diagnostics were used to assay for total cholesterol and triglycerides.

### Chromatin immunoprecipitation

ChIP was performed using the MAGnify Chromatin Immunoprecipitation System (Thermo Fisher Scientific) according to the manufacturer’s instruction. Frozen liver tissue (50mg) was minced in DPBS and homogenized with a dounce tissue grinder. At room temperature, homogenized tissue was crosslinked with 1% formaldehyde for 10 minutes then quenched with 0.125 M glycine for 5 minutes, swirling gently about every 2 minutes during incubation. The crosslinked mixture was centrifuged at 200 × g for 10 minutes at 4°C then washed twice with cold PBS buffer and resuspended in lysis buffer supplemented with proteinase inhibitors for 1 hour at 4°C. The lysates were sonicated with a Covaris E220 (Peak incident power – 140 Watts, Duty factor – 5%, Cycle/bust – 200, Duration – 75 seconds) to yield 200-500 base pair DNA fragments. Samples were centrifuged at 12,000 × g for 15 minutes at 4°C to pellet debris and the chromatin-containing supernatant was transferred to new sterile tubes.

Next, the chromatin samples were diluted in cold dilution buffer with protease inhibitors to a final volume of 100 μl per reaction and 10 μl from each chromatin sample was reserved for input control. Coupled antibody-Dynabeads were prepared by adding Dynabeads protein A/G and 2 μg antibody [H3K9ac (Active Motif) or H3K27ac (Abcam)] to 100 μl ice-cold dilution buffer and rotating the tubes end-over-end at 4°C for 1 hour. IP was performed by incubating diluted chromatin with the appropriate antibody-Dynabeads complex and rotating the tubes end-over-end at 4°C for 2 hours. Using a DynaMag-PCR magnet (Thermo Fisher Scientific), the beads were washed 3 times each with ice-cold IP buffer 1 and twice with ice-cold IP buffer 2. The supernatant was discarded, and reverse cross-linking buffer prepared with proteinase K was added into the IP sample tubes to reverse the cross-linking. The beads were fully resuspended by vortexing and incubated at 55°C for 15 minutes in a C1000 Touch thermal cycler (BIO-RAD) then re-pelleted with the magnet. The IP sample liquids were transferred to new sterile tubes, incubated at 65°C for 15 minutes, and cooled on ice for 5 minutes.

To purify the DNA, magnetic purification beads prepared with DNA purification buffer were added to each tube. After pipetting up and down 5 times to mix, samples were incubated at room temperature for 5 minutes, beads were pelleted with the magnet, and the supernatant discarded. The beads were washed 2 times using DNA wash buffer following the same procedure. The DNA was eluted with DNA elution buffer by incubating at 55°C for 20 minutes in a thermal cycler. The purified DNA was analyzed by ChIP-qPCR data relative to input using the mouse negative control and mouse positive control primer set Gapdh in the ChIP-IT qPCR analysis kit (Active Motif). Reaction conditions were programmed with initial denaturation at 50°C for 2 minutes and 95°C for 10 minutes followed by 40 cycles of 95°C for 15 seconds and 60°C for 1 minute. Relative percent of input enrichment and fold enrichment were calculated with the formulas (100*2^ (Cq Adjusted input-Cq IP)) and 2-ΔΔCq respectively.

### ChIP-sequencing analysis

KAPA Hyper Prep Kit (Kapa Biosystems) was used for end repair, A-tailing, and adapter ligation with the KAPA Unique Dual-Indexed Adapter Kit (Kapa Biosystems) according to the manufacturer’s instructions. Briefly, ChIP DNA was subjected to end repair and A-tailing by adding End Repair & A-Tailing buffer and enzyme mix, incubating at 20°C for 30 minutes followed by 60°C for 30 minutes, then cooling to room temperature. Adaptor ligation was performed by adding adapter stock, ligation buffer, DNA ligase, and PCR-grade water to the reaction product, and incubating at 20°C for 25 minutes. Post-ligation clean-up was performed with 0.8X KAPA Pure Beads (Kapa Biosystems); beads were mixed with the ligation product and incubated for 15 mins to bind DNA to the beads. The bead/DNA complex was captured using a DynaMag-PCR magnet (Thermo Fisher Scientific) and washed twice with 80% ethanol. Purified DNA was eluted with DNA elution buffer at room temperature for 2 minutes. Library amplification was performed with KAPA HiFi HotStart ReadyMix and Library Amplification Primer Mix and the following PCR program: initial denaturation 98°C for 45 seconds, denaturation 98°C for 15 seconds, annealing 60°C for 30 seconds, extension 72°C for 30 seconds, and final extension 72°C for 1 minute then hold 4°C. Library amplification was carried out for 7 (H3K27ac) and 13 cycles (H3K9ac) plus 1 full cycle on the PCR product to get rid of PCR bubbles. A 0.7X-0.9X KAPA Pure Beads double-sided size selection was done after PCR amplification. An Agilent Bioanalyzer 2100 was used to profile the sizes, concentrations, and qualities of the libraries. Libraries were subjected to high-throughput sequencing by Novogene Corporation, Inc. on an Illumina NovaSeq S4 platform and 150-bp paired-end reads were generated.

ChIP-seq data analysis was performed by the UC Davis Bioinformatics Core. The raw read data was filtered using HTStream (version 1.2.0) which included screening for contaminants (such as PhiX), removing PCR duplicated reads, overlapping paired reads, quality-based trimming, and adapter trimming. BWA MEM (version 0.7.16a) was used to align the processed data to the mouse genome (GRCm38). Calling peaks was done using phantompeakqualtools (version 1.2.2). Bioconductor (version 3.11) package DiffBind (version 2.12.0) was used to combine raw count data for all peaks across replicates using R (version 3.6.3). Finally, GREAT (version 4.0.4) was used to find the regulatory domains for the peaks and the genes within those domains.

### RNA-sequencing analysis

RNA-seq library production and sequencing were performed by Novogene Corporation, Inc. RNA degradation and contamination were monitored on 1% agarose gels and purity was checked using the NanoPhotometer spectrophotometer (Implen GmbH). RNA integrity and quantitation were assessed using the RNA Nano 6000 Assay Kit of the Agilent Bioanalyzer 2100 system. A total of 1 μg RNA per sample was used as input material. Sequencing libraries were generated using NEBNext Ultra RNA Library Prep Kit for Illumina (New England Biolabs) following manufacturer’s recommendations and index codes were added to attribute sequences to each sample. Briefly, mRNA was purified from total RNA using poly-T oligo-attached magnetic beads. Fragmentation was carried out using divalent cations under elevated temperature in NEBNext First Strand Synthesis Reaction Buffer. First-strand cDNA was synthesized using random hexamer primers and M-MuLV Reverse Transcriptase (RNase H-). Second-strand cDNA synthesis was subsequently performed using DNA Polymerase I and RNase H. Remaining overhangs were converted into blunt ends via exonuclease/polymerase activities. After adenylation of 3’ ends of DNA fragments, NEBNext adaptors with hairpin loop structure were ligated to prepare for hybridization. In order to preferentially select cDNA fragments of ∼150-200 bp in length, the library fragments were purified with the AMPure XP system (Beckman Coulter). Three μl USER Enzyme (New England Biolabs) was used with size-selected, adaptor-ligated cDNA at 37°C for 15 minutes followed by 5 minutes at 95°C before PCR. PCR was performed with Phusion High-Fidelity DNA polymerase, NEBNext Universal PCR primer for Illumina, and NEBNext Index primer (New England Biolabs). PCR products were purified again with the AMPure XP system and library quality was assessed on the Agilent Bioanalyzer 2100. The clustering of the index-coded samples was performed on a cBot Cluster Generation System using PE Cluster Kit cBot-HS (Illumina) according to the manufacturer’s instructions. After cluster generation, the library preparations were sequenced on an Illumina NovaSeq S4 platform and 150-bp paired-end reads were generated.

Raw reads (FASTQ) were processed through fastp (0.20.0), removing reads containing adapter and poly-N sequences and low-quality reads while simultaneously calculating Q20, Q30, and GC content. Paired-end clean reads were aligned to the mouse reference genome (grcm38) using the Spliced Transcripts Alignment to a Reference (STAR) software (v2.6.1d). FeatureCounts (v1.5.0-p3) was used to count the reads mapped to each gene; RPKM of each gene was calculated based on the length of the gene and reads mapped to the gene.

### Statistical Analysis

ChIP-seq: Bioconductor packages edgeR (version 3.26.8) and limma (version 3.40.6) were used to perform differential expression analysis across the called peaks. Data were prepared by first choosing to keep peaks that achieved at least 1 count per million (cpm) in at least 1 sample; normalization factors were calculated using trimmed mean of M-value (TMM); and the voom transformation was applied. A completely randomized design was implemented, comparisons of interest were extracted using contrasts, and moderated statistics were computed using the empirical bayes procedure eBayes. Finally, each peak was corrected for multiple testing using the Benjamini-Hochberg false discovery rate correction. Differential expression was defined as an adjusted p-value <0.05.

RNA-seq: Differential expression analysis between two groups (three biological replicates per group) was performed using DESeq2 R package (v1.20.0). The resulting p-values were adjusted using the Benjamini and Hochberg approach for controlling the false discovery rate. Genes with an adjusted p-value <0.05 found by DESeq2 were considered differentially expressed.

qPCR and Western blot: Student’s t-test was used to compare differences between two groups. For multiple-group comparisons, Kruskal-Wallis one-way ANOVA followed by uncorrected Dunn’s test was performed with GraphPad Prism software (version 9.0). Data are presented as mean ± SEM, and p<0.05 was considered statistically significant.

### Other

InteractiVenn (http://www.interactivenn.net/index.html), a web-based tool, was used for identifying differentially expressed co-upregulated and co-downregulated genes between RNA- and ChIP-seq analysis and creating Venn diagrams. Gene ontology enrichment analyses of differentially expressed genes were conducted with the web-based server, g:Profiler (https://biit.cs.ut.ee/gprofiler/gost). LISA (epigenetic Landscape In Silico deletion Analysis, lisa.cistrome.org) was used to determine functional relevance of differentially expressed TF genes.

## Acknowledgments

The research was supported by the National Institutes of Health through grant numbers R01DK104770 (to V.M.), R01ES029213 and R01AA027075 (to J.M.L), R01HL113178 (to E.A.G), and DK092993 (to UCD Mouse Metabolic Phenotyping Center).

## Abbreviations

“ac” suffix: acetylated
AICAR: 5-aminoimidazole-4-carboxamide-1-β-D-ribofuranoside
AMPK: AMP-activated protein kinase
C3H: C3HeB/FeJ
ChIP: chromatin immunoprecipitation
HDAC: histone deacetylase
“me3” suffix: trimethylated
“p” prefix: phosphorylated
PCA: D-penicillamine
PPARα: peroxisome proliferator-activated receptor alpha
PPARγ: peroxisome proliferator-activated receptor gamma
SAH: S-adenosylhomocysteine
TBST: tris-buffered saline with Tween-20
TF: transcription factor
tx-j: Jackson Laboratory toxic milk mouse C3He-Atp7btx-J/J
WD: Wilson disease.

## Data deposition

The data discussed in this publication have been deposited in NCBI’s Gene Expression Omnibus (Edgar et al., 2002) and are accessible through GEO Series accession number GSE168972 (https://www.ncbi.nlm.nih.gov/geo/query/acc.cgi?acc=GSE168972).

