## Supplementary data 1 for "Wilson disease: intersecting DNA methylation and histone acetylation regulation of gene expression in a mouse model of hepatic copper accumulation"

**Table of Contents**

| Section | Page |
| --- | --- |
| Supplementary Figures | 2 |
| Supplementary Tables | 9 |

**Figure S1: Animal study design.**

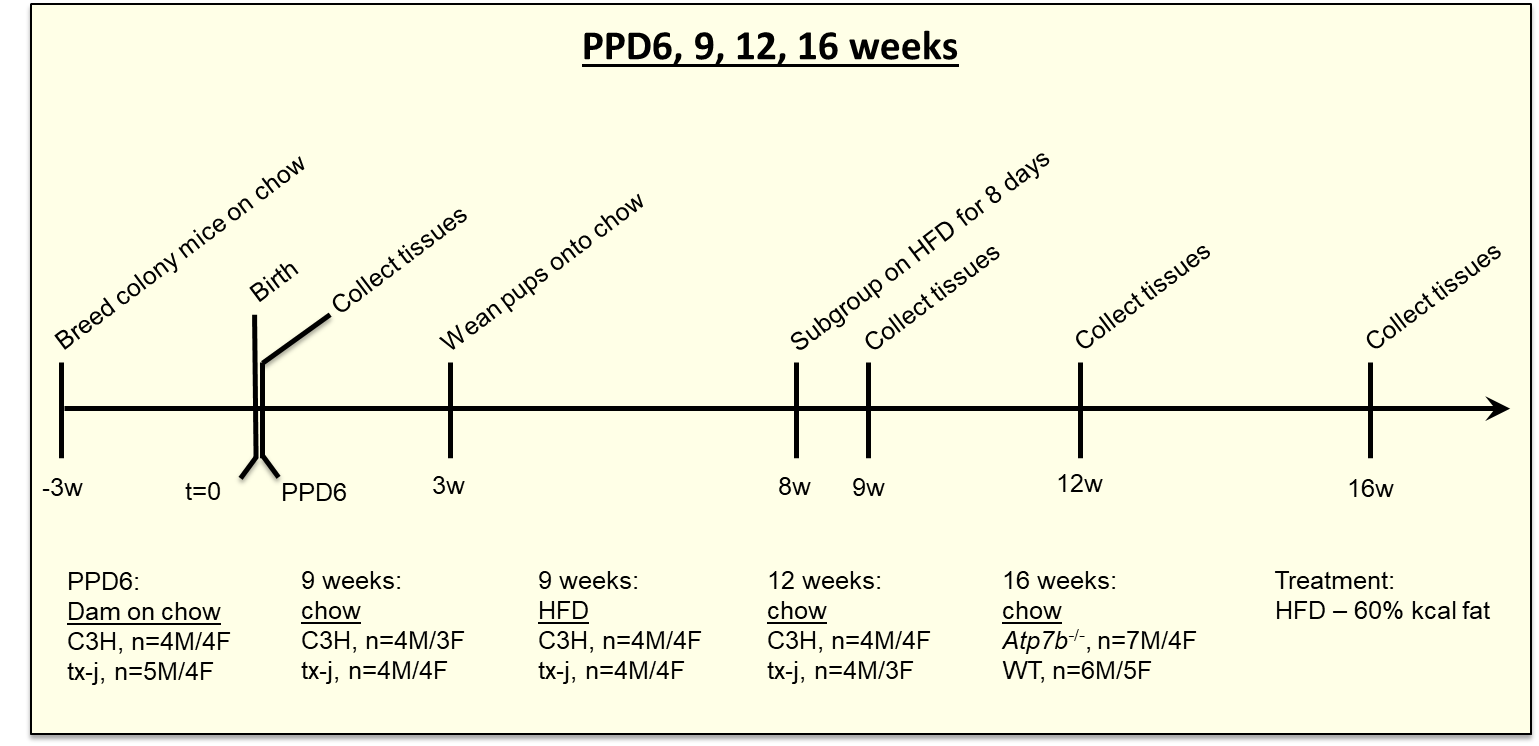

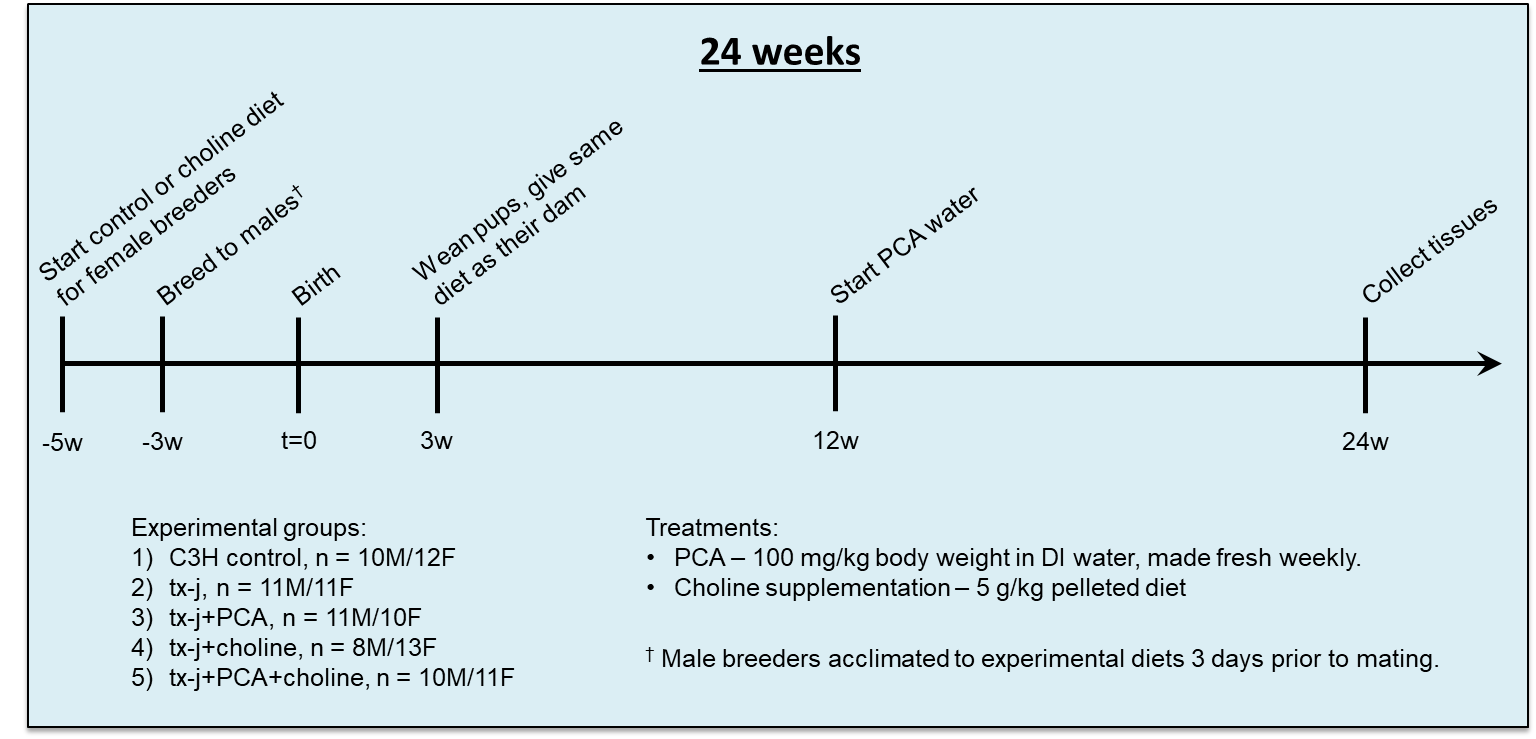

C3H, C3HeB/FeJ; HFD, high fat diet; PCA, D-penicillamine; PPD6, post-partum day 6; tx-j, C3He-Atp7b^tx-j^/J; WT, wild-type.

**Figure S2: Age- and dose-dependent immunoblots of HDAC4 and HDAC5 in mouse and HepG2 models of WD.**

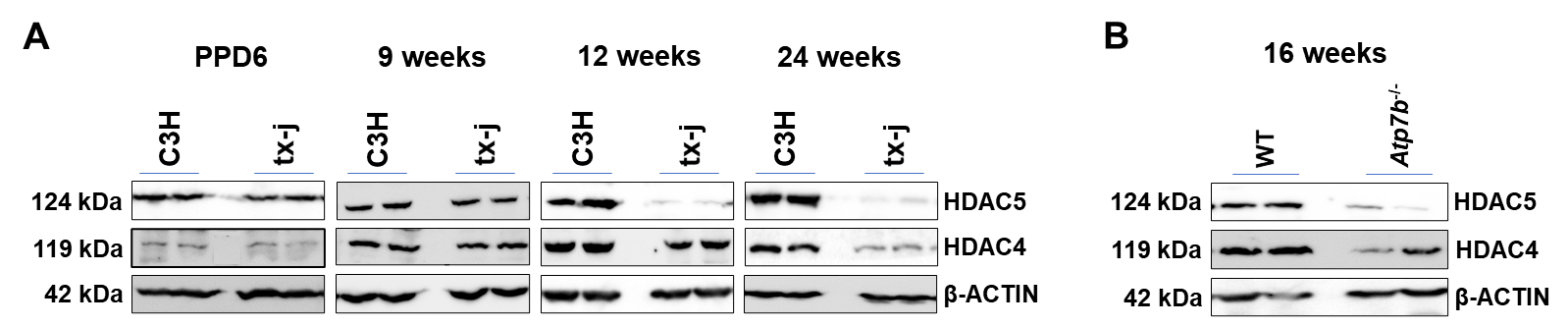

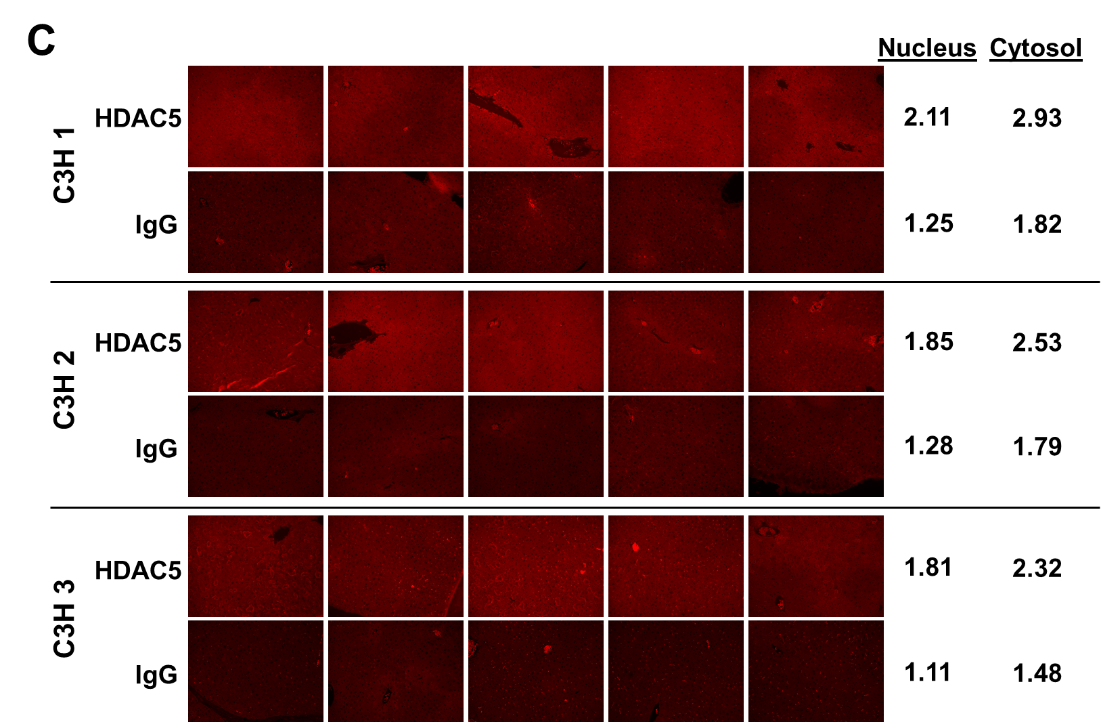

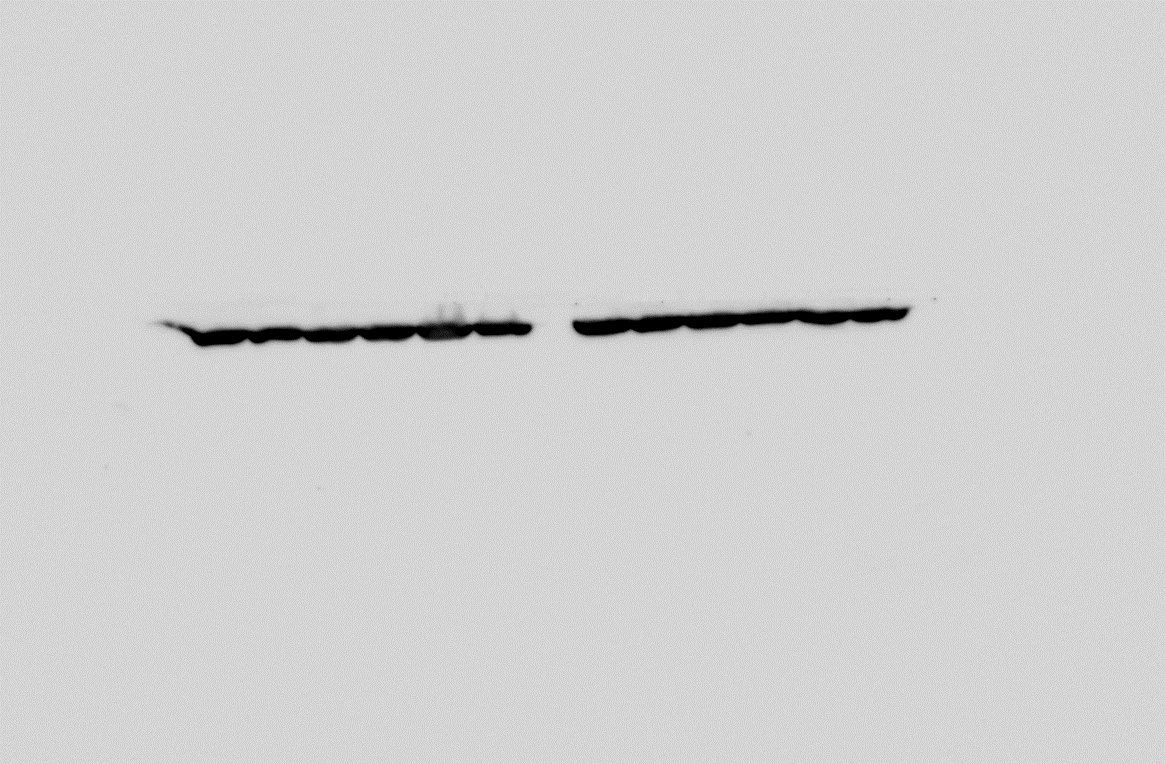

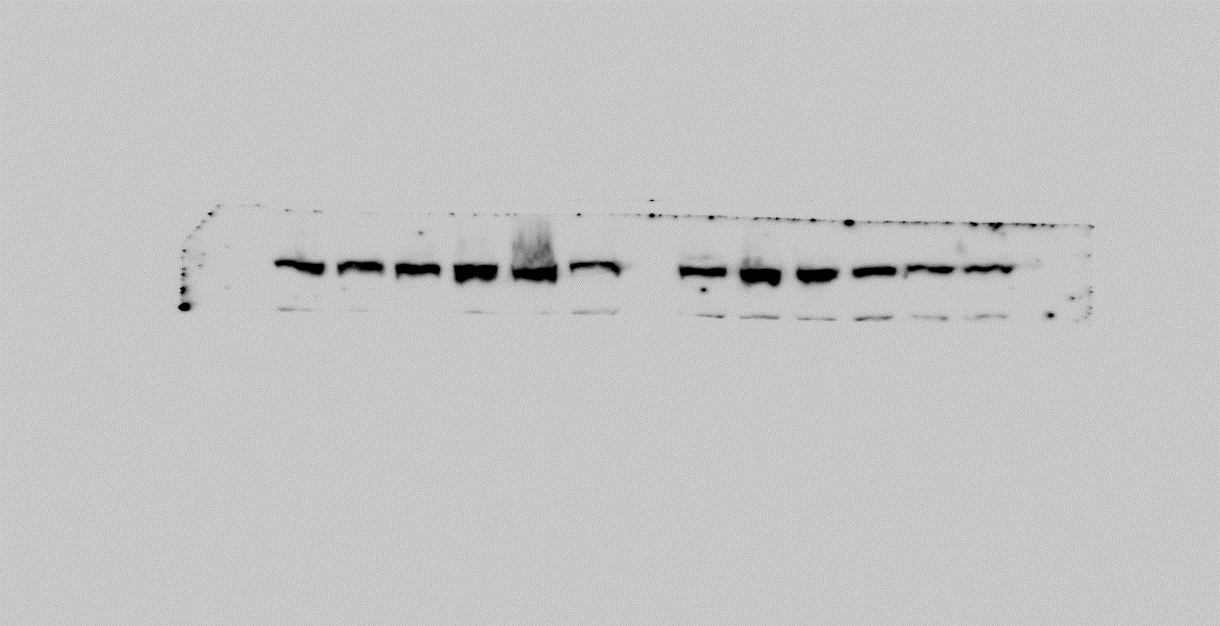

**42 kDa**

**124 kDa**

**HDAC5**

**β-ACTIN**

**µM	0	5	10	25**

**HepG2 cells: IPN treatment**

**D**

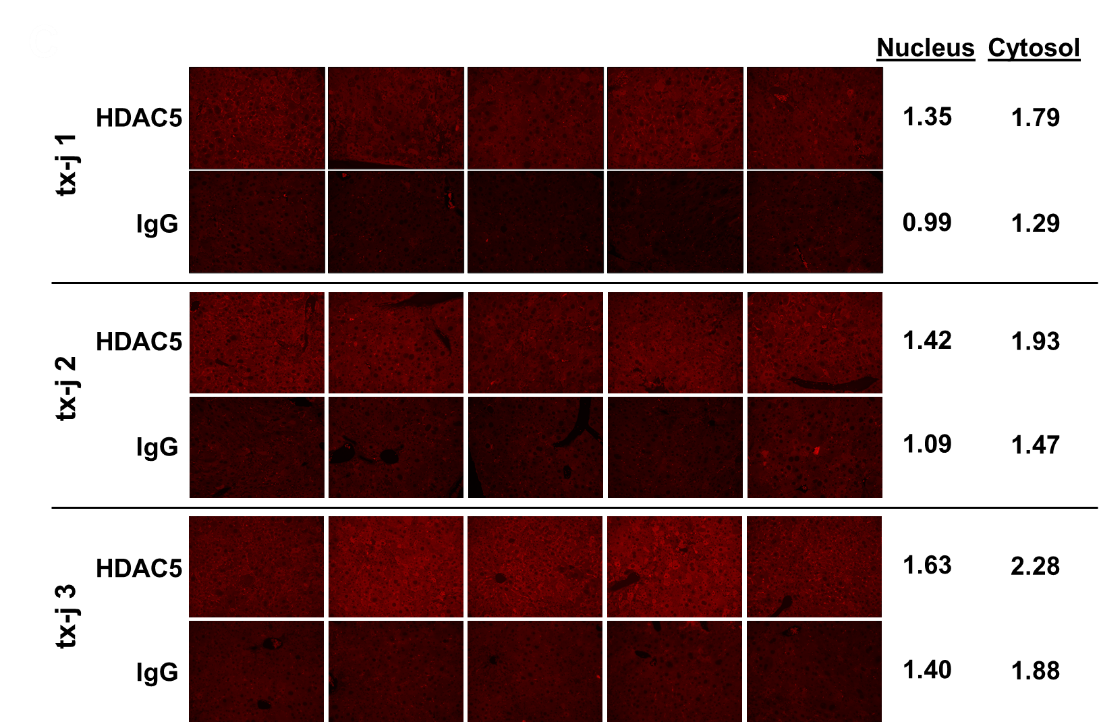

A: Total protein liver lysate for histone deacetylase (HDAC) 4 and HDAC5 protein expression in tx-j mice compared to C3H control at post-partum day 6 (PPD6; C3H n=4M/4F, tx-j n=5M/4F), 9 weeks (C3H n=4M/3F, tx-j n=4M/4F), 12 weeks (C3H n=4M/4F, tx-j n=4M/3F), and 24 weeks (C3H n=10M/12F, tx-j n=11M/11F). Immunoblot images show 2 representative samples per group per time point. B: Total protein liver lysate for HDAC4 and HDAC5 protein expression in 16-week old *Atp7b*^-/-^ (n=7M/4F) mice and wild-type (WT, n=6M/5F). Immunoblot images show 2 representative samples per group. C: Immunoblot of HepG2 cell lysates treated with 50µM of CuSO_4_ for 24 hours followed by 24-hour treatment with isoproterenol (IPN, 0-25µM, n=3), a HDAC5 activator. β-ACTIN was used as loading control in all immunoblots.

**Figure S3: Immunoblots of AMPKα signaling in 24-week old tx-j mice and AICAR treatment in HepG2 cells.**

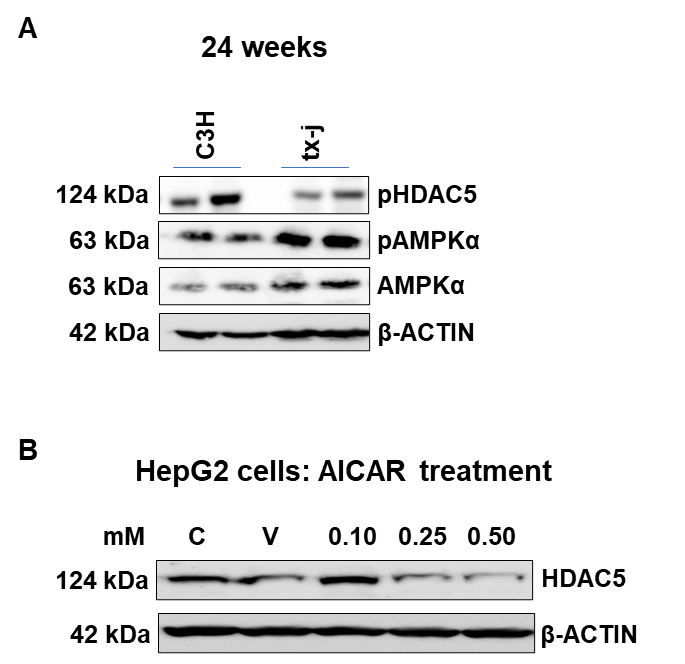

A: Total protein liver lysates of total AMPKα, phosphorylated AMPKα (pAMPKα), and phosphorylated HDAC5 (pHDAC5) obtained from 24-week old C3H (n=3M/3F) and tx-j (n=3M/2F). Immunoblot images show 2 representative samples per group. B: HDAC5 immunoblot of HepG2 cell lysates treated with 50 µM CuSO_4_ for 24 hours followed by 5-aminoimidazole-4-carboxamide-1-β-D-ribofuranoside (AICAR) treatment (0-0.5 mM; n=3), an AMPK activator, for 24 hours. C=control, V=treated with vehicle (DMSO) only.

**Figure S4: Immunoblots of HDACs and H3 marks, and *Hat1* transcript levels, in response to PCA and choline.**

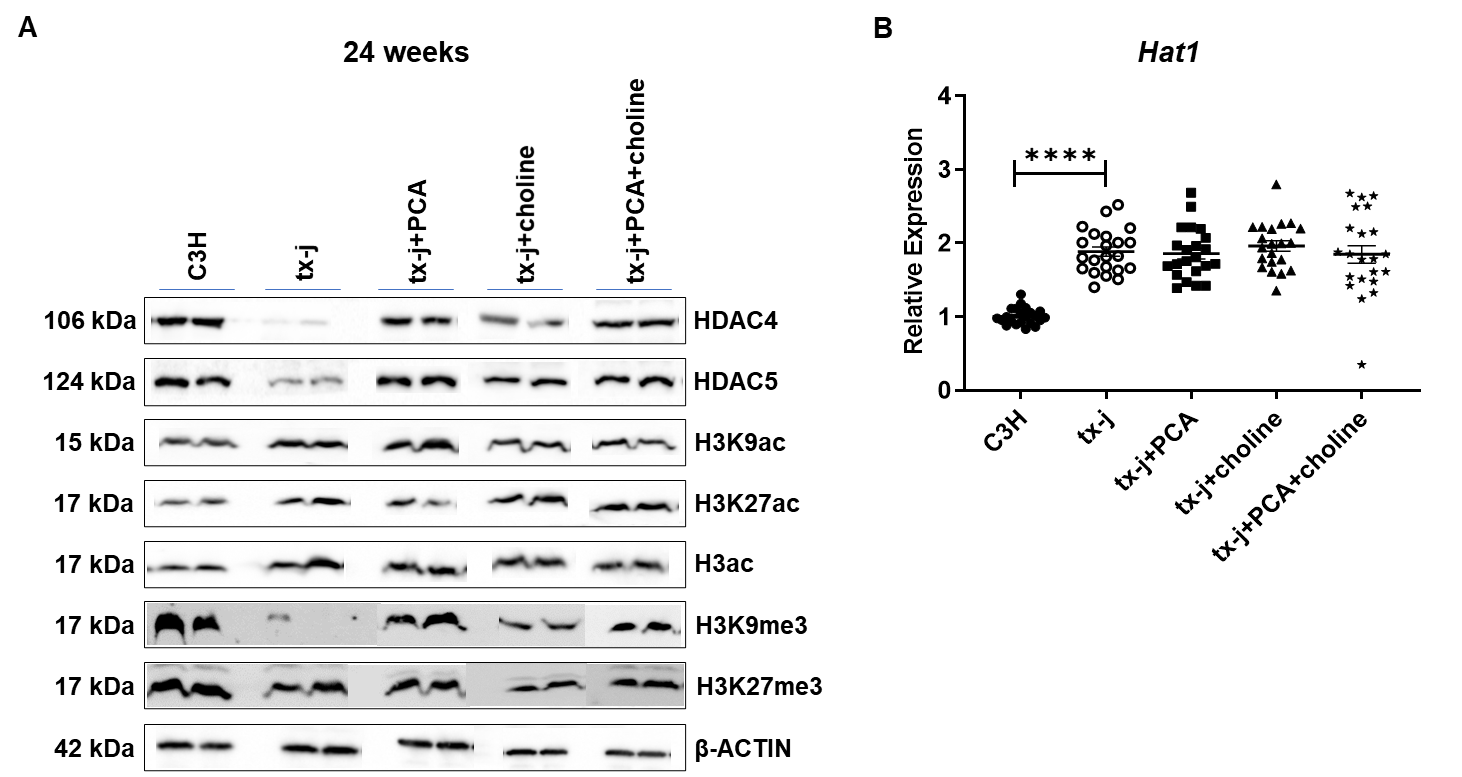

A: Immunoblots of total protein liver lysates from 24-week old C3H (n=10M/12F) and tx-j mice (n=11M/11F), and tx-j mice treated with PCA (n=11M/10F), choline (n=8M/13F), and PCA+choline (n=10M/11F). Immunoblot images show 2 representative samples per group. B: Transcript levels of histone acetyltransferase (*Hat1*) normalized to *Gapdh*. Data are represented as means ± SEM and statistical significance was determined by Kruskal-Wallis one-way ANOVA followed by uncorrected Dunn’s test (****p< 0.001).

**Figure S5: ChIP-seq and RNA-seq volcano plots.**

Volcano plots showing the separation of differentially regulated genes (p<0.05) in both ChIP-seq (left) and RNA-seq (right) comparing livers of 3M/3F 24-week old C3H vs. tx-j mice. Data are represented as -log p-value vs. log2 fold change.

**Figure S6: Transcription factor enrichments.**

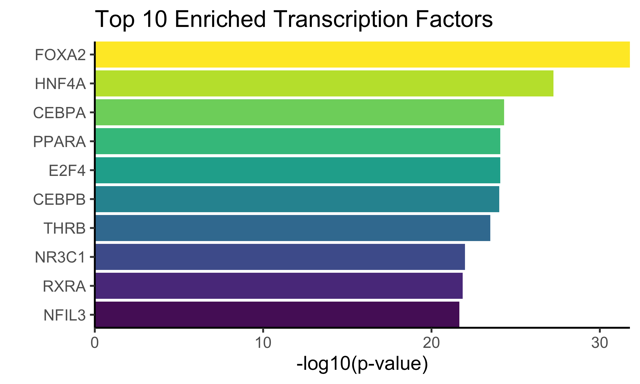

LISA (epigenetic Landscape In Silico deletion Analysis) from Cistrome.org, which utilizes publicly available ChIP-seq data for transcription factors and chromatin regulators, was used to determine gene enriched transcription factors. Bar graph shows top 10 TFs enriched with differentially expressed co-upregulated and co-downregulated genes between ChIP- and RNA-seq.

**Figure S7: Immunoblots of HDAC5, PPARγ, and PPARα, and transcript levels of *Hmox1*.**

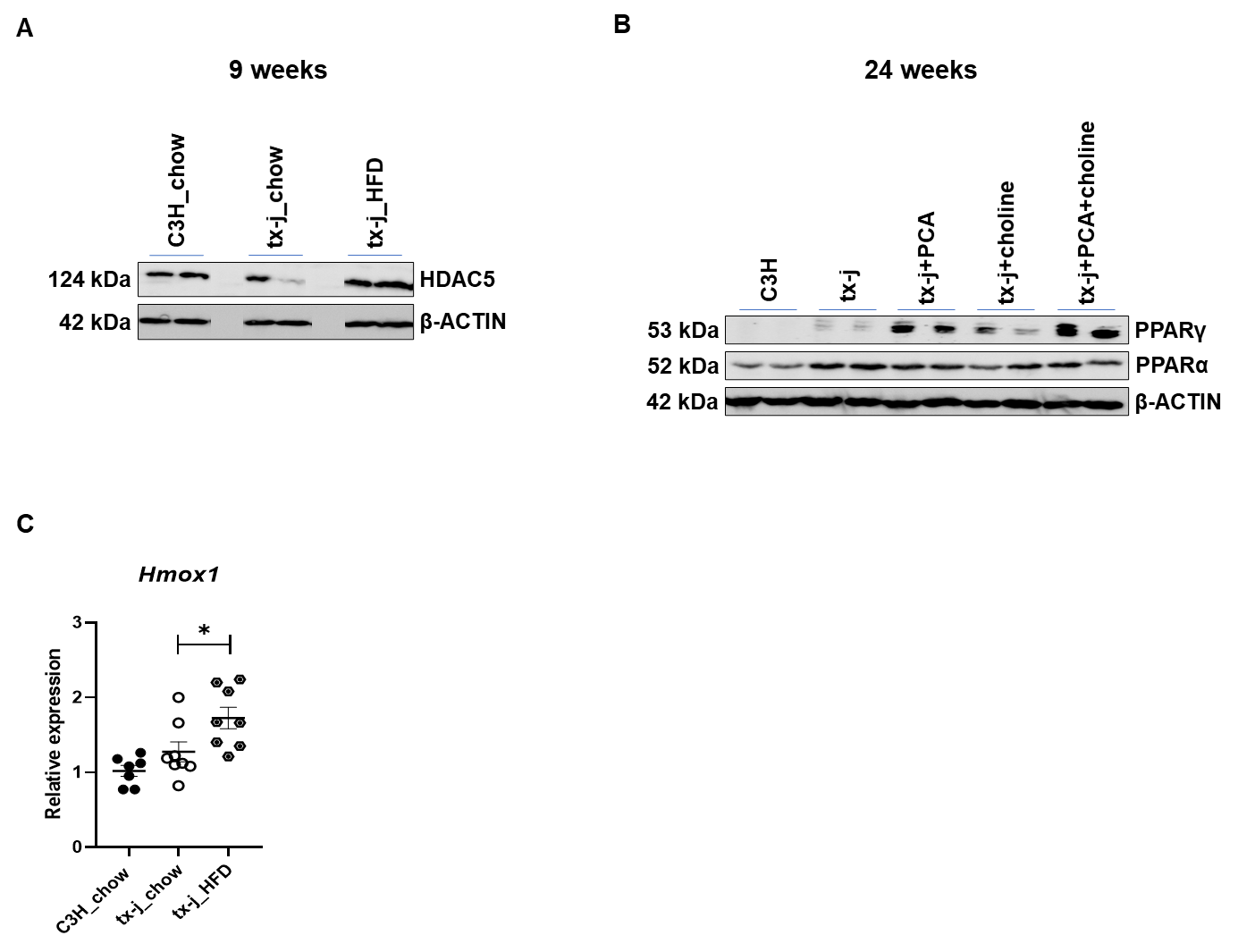

A: HDAC5 protein expression in total protein liver lysate from 9-week mice on chow (C3H n=4M/3F, tx-j n=4M/4F) and fed a 60% kcal fat diet (HFD, n=4M/4F each) for 8 days. B: Immunoblots of PPARγ and PPARα in total protein liver lysate of 24-week old C3H, tx-j, tx-j+PCA, tx-j+choline, and tx-j+PCA+choline (n=3M/3F per group). Immunoblot images show 2 representative samples per group. C: Liver transcript levels of *Homx1* are normalized to *Gapdh* in 9-week mice on chow or HFD. Data are represented as means ± SEM and statistical significance was determined by Kruskal-Wallis one-way ANOVA followed by uncorrected Dunn’s test (*p< 0.05).

**Table S13: Primers for qPCR analyses.**

| *Gene* | Full name | Primer | Sequence 5' to 3' | Accession | Exon-exon overlap | % Primer efficiency |
| --- | --- | --- | --- | --- | --- | --- |
| *Aebp1* | AE binding protein1 | F | CCTCGCTCTGGGACTTTCAA | NM_009636.3 | No | 99.47 |
|  |  | R | CGGGGTAGCTCACTCTCGTG |  | Yes |  |
| *Aebp2* | AE binding protein2 | F | AGCTCGCCATGTACCCACAC | NM_001005605.2 | No | 94.76 |
|  |  | R | CCTCCGCTTGTTCATTCCAG |  | No |  |
| *Ampka1* | Protein kinase, AMP-activated, alpha 1 catalytic subunit | F | GAACGCATTTGGAGGACATGA | NM_001013367 | No | 99.35 |
|  |  | R | AGCCCCGAACAAAAAGAAGC |  | No |  |
| *Atf4* | Activating transcription factor 4 | F | GGAAGCCTGACTCTGCTGCT | NM_001287180.1 | No | 95.64 |
|  |  | R | CTCCGGGCTCATACAGATGC |  | No |  |
| *Bcl2* | B cell leukemia/lymphoma 2 | F | AGGGCCTGAACTTGCGTGAA | NM_009741.5 | No | 102 |
|  |  | R | AGACAAGCGAGCTGGACAGG |  | No |  |
| *Cd36* | CD36 molecule | F | TGGGACCATTGGTGATGAAA | NM_001159555 | No | 99.65 |
|  |  | R | CCAATCCCAAGTAAGGCCATC |  | No |  |
| *Chrebp* | Carbohydrate response element binding protein (ChREBP | F | CCCCTCAGACACCCACATCTTC | NM_021455.5 | Yes | 95.2 |
|  |  | R | AGACGAGGCTGGAGATGGCA |  | No |  |
| *Fxr* | Farnesoid X receptor | F | CGAATGCCCGCTGAGACTGG | NM_009108.2 | No | 94.23 |
|  |  | R | CCCCTTTTATTCTGCCTGCCGA |  | No |  |
| *Gapdh* | Glyceraldehyde-3-phosphate dehydrogenase | F | GAAGCTTGTCATCAACGGGAAG | NM_008084.2 | No | 102.1 |
|  |  | R | TTTGATGTTAGTGGGGTCTCGC |  | No |  |
| *Gpx3* | Glutathione peroxidase 3 | F | TGGCTTGGTCATTCTGGGCTT | NM_008161.4 | No | 99.64 |
|  |  | R | CCCCACCTGGTCGAACATACTT |  | Yes |  |
| *Hat1* | Histone acetyltransferase 1 | F | CAGCGGAAGATCCGTCCA | NM_026115 | No | 106.2 |
|  |  | R | GCCATATCTTCACTGAATCCTTGC |  | Yes |  |
| *Hmgb2* | HMG box domain containing 4 | F | AGGAGACAGAGGGCATGGTG | NM_178017.2 | Yes | 107.32 |
|  |  | R | GGACAGCCGTTCAGTTCAGG |  | No |  |
| *Hmox1* | Heme oxygenase 1 | F | ATGCCCCAGGATTTGTCTGA | NM_010442.2 | No | 96.99 |
|  |  | R | TCTGGACACCTGACCCTTCTG |  | No |  |
| *Hnf4a* | Hepatic nuclear factor 4, alpha | F | CGCCCCGGTTGACTCTTGAT | NM_008261.3 | No | 96.4 |
|  |  | R | TCCAGTCTCACAGCCCATTCCT |  | No |  |
| *Hnf6* | Hepatic nuclear factor 4 | F | GGGTTGGAGCTGAGCACTGT | NM_008262.3 | No | 99.62 |
|  |  | R | TGCTCGATGAGGACGATGAA |  | No |  |
| *Irf6* | Interferon regulatory factor 6 | F | GCTCATTGCCCACCAGAAAG | NM_016851.2 | No | 98.64 |
|  |  | R | ACTGGGATGACCTGGACCAA |  | Yes |  |
| *Klf4* | Kruppel-like factor 4 | F | CACAGGCGAGAAACCTTACCA | NM_010637.3 | Yes | 97.12 |
|  |  | R | CGCACTTCTGGCACTGAAAG |  | No |  |
| *Lpl* | lipoprotein lipase | F | AACCAAGTCTGGCCTCGAACT | NM_008509 | No | 101.2 |
|  |  | R | AAGCTCCCAGGACACAGGAA |  | No |  |
| *Mmp12* | matrix metallopeptidase 12 | F | TGGAATATGACCCCCTGTTC | NM_008605.3 | No | 105.2 |
|  |  | R | CCAGCAAGCACCCTTCACTA |  | No |  |
| *Pparα* | peroxisome proliferator-activated receptor alpha | F | CGATGCTGTCCTCCTTGATGA | NM_011144.6 | No | 98.5 |
|  |  | R | GAAGTCAAACTTGGGTTCCATGAT |  | No |  |
| *Pparγ* | Peroxisome proliferator activated receptor gamma | F | CCGAAGAACCATCCGATTGA | NM_011146.3 | No | 105.27 |
|  |  | R | GAGACATCCCCACAGCAAGG |  | No |  |
| *Smad4* | SMAD family member 4 | F | TTGCTGAGTCTTCCCTGCTGG | NM_008540.3 | No | 98.63 |
|  |  | R | AGCTCTGACTCTCCTCGGCA |  | No |  |
| *Sod2* | Superoxide dismutase 2, mitochondrial | F | CAAGCACAGCCTCCCAGACC | NM_013671.3 | No | 98.08 |
|  |  | R | TGGCGTTGAGGTTGTTCACGT |  | No |  |
| *Sox6* | SRY (sex determining region Y)-box 6 | F | GCGACACAATGTACTGCGTTTT | NM_009238.3 | No | 97.09 |
|  |  | R | TGCTATCACCATGCCATAGGAC |  | No |  |
| *Srebf1* | sterol regulatory element binding transcription factor 1 | F | CTGGCTTGGTGATGCTATGTTG | NM_011480.3 | No | 104.9 |
|  |  | R | GACCATCAAGGCCCCTCAA |  | No |  |
| *Stard6* | StAR-related lipid transfer (START) domain containing 6 | F | AAAATTGCCTGCATCGGTAA | NM_029019.4 | No | 104.07 |
|  |  | R | CACGTATGGGTGGAATTCTG |  | Yes |  |
| *Ucp2* | Uncoupling protein 2 (mitochondrial, proton carrier | F | CGTCATCGCCTCCCCTGTTG | NM_011671.5 | No | 98.88 |
|  |  | R | CCGGAGCATGGTAAGGGCAC |  | No |  |

**Table S14: Antibodies for immunoblotting.**

| **PRIMARY ANTIBODIES** | | | | |
| --- | --- | --- | --- | --- |
| **Antibody** | **MW** | **Vendor** | **Cat. No.** | **Dilution** |
| AMPKα | 62 | Cell Signaling Technology | 23A3 | 1:1000 |
| H3ac | 17 | Millipore | 06-599 | 1:500 |
| H3K27ac | 15 | Abcam | ab177178 | 1:1000 |
| H3K27me3 | 15 | Sigma-Aldrich | 07-449 | 1:500 |
| H3K9ac | 17 | PMT BIOLABS | PMT-156 | 1:2000 |
| H3K9ac | 17 | Active Motif | 39918 | 1:500 |
| H3K9me3 | 17 | Abcam | ab8898 | 1:1000 |
| HDAC5 | 124 | Cohesion Biosciences | CPA2445 | 1:500 |
| HDAC4 | 119 | Boster Biological Technology | A00971-1 | 1:500 |
| PPARα | 52 | Abcam | ab24509 | 1:500 |
| PPARγ | 53 | Cell Signaling Technology | 81B8 | 1:1000 |
| pAMPKα | 63 | Cell Signaling Technology | 2535 | 1:1000 |
| pHDAC5 | 124 | Affinity Biosciences | ABIN6267700 | 1:500 |
| β -ACTIN | 42 | Sigma-Aldrich | A5441 | 1:5000 |
| **SECONDARY ANTIBODIES** | | | | |
| **Antibody** |  | **Vendor** | **Cat. No.** | **Dilution** |
| Anti-Mouse |  | Jackson Immunoresearch Laboratoreis | 115-035-003 | 1:10000 |
| Anti-Rabbit |  | Jackson Immunoresearch Laboratories | 111-035003 | 1:10000 |
